# STAPLE: automating spatial transcriptomics analysis and AI interpretation

**DOI:** 10.64898/2026.03.30.715127

**Authors:** Dmitrijs Lvovs, Jeffrey Quinn, André Forjaz, Ivette Santana-Cruz, Orian Stapleton, Kranthi Vavikolanu, Meredith Wetzel, Demystifying Pancreatic Cancer Therapies TeamLab, Data Science Hub Teamlab, Vincent Bernard, Brian R. Herb, Alexander Favorov, Luciane T. Kagohara, Ashley L Kiemen, Anirban Maitra, Dimitrios N. Sidiropoulos, Wesley Tansey, Laura D Wood, Atul Deshpande, Michael Noble, Elana J. Fertig

## Abstract

Spatial transcriptomics workflows often span separate tools for cell typing, neighborhoods, and cell–cell communication, yielding fragmented outputs that hinder scalability, interpretation, and reproducibility. STAPLE systematizes analyses across distinct methods into a modular framework, unifying data structures and cross-tool interoperability. End-to-end analyses are performed unassisted with a single invocation, fostering rigorous, reproducible spatial transcriptomics analysis. Its novel, AI-enabled reporting layer synthesizes quantitative results into summaries of biological findings, facilitating analysis interpretation.

## Main

Spatial transcriptomics (ST) measures gene expression directly in tissues, enabling comprehensive characterization of molecular and cellular changes resulting from distinct cellular niches to be inferred in a single assay. Such multi-scale inference requires applying multiple bioinformatics tools in series, often manually, into a comprehensive analysis^1^. Fragmentation of bioinformatics analysis pipelines makes it challenging to connect results across software modules and is exacerbated when tools vary in the data formats supported, their language of implementation or method of operation. Different tools have distinct inputs, and disregard clinical metadata or comparative analysis when focused on single-sample cellular and molecular interpretation. Each analysis module also outputs ranked lists containing multiple molecular and cellular features inferred from the data, that require intensive expert curation and manual coding for cross-sample comparisons. Together, these factors hamper efficiency, scalability, reproducibility, and interpretability.

We developed STAPLE: the Spatial Transcriptomics Analysis Pipeline (Figure 1) to address these challenges and systematize end-to-end ST analysis. Combining open-source bioinformatics software in a modular framework with bespoke tooling, guided by flexible configuration, STAPLE automatically computes cellular annotations, spatial statistics, and cell-to-cell communication. STAPLE execution is orchestrated with Nextlow^2^ and operates in 5 phases (Fig. 1a): data ingest, preprocessing, cell type annotation, ligand-receptor and spatial analysis, and reporting. Its modular design enables runtime workflow steering via parameters that select specific sub-tools for analysis tasks. Results for each analysis (Fig. 1b) are automatically summarized into standard figures and tables (Fig. 1c). To our knowledge, STAPLE is the first ST pipeline to incorporate comprehensive, AI-enabled interpretation of ST analysis. This AI-integration links cellular and molecular features to biological and clinical interpretations based on literature, applicable for the findings in individual samples, comparisons between sample groups, and metadata of the samples.

**Fig. 1.**
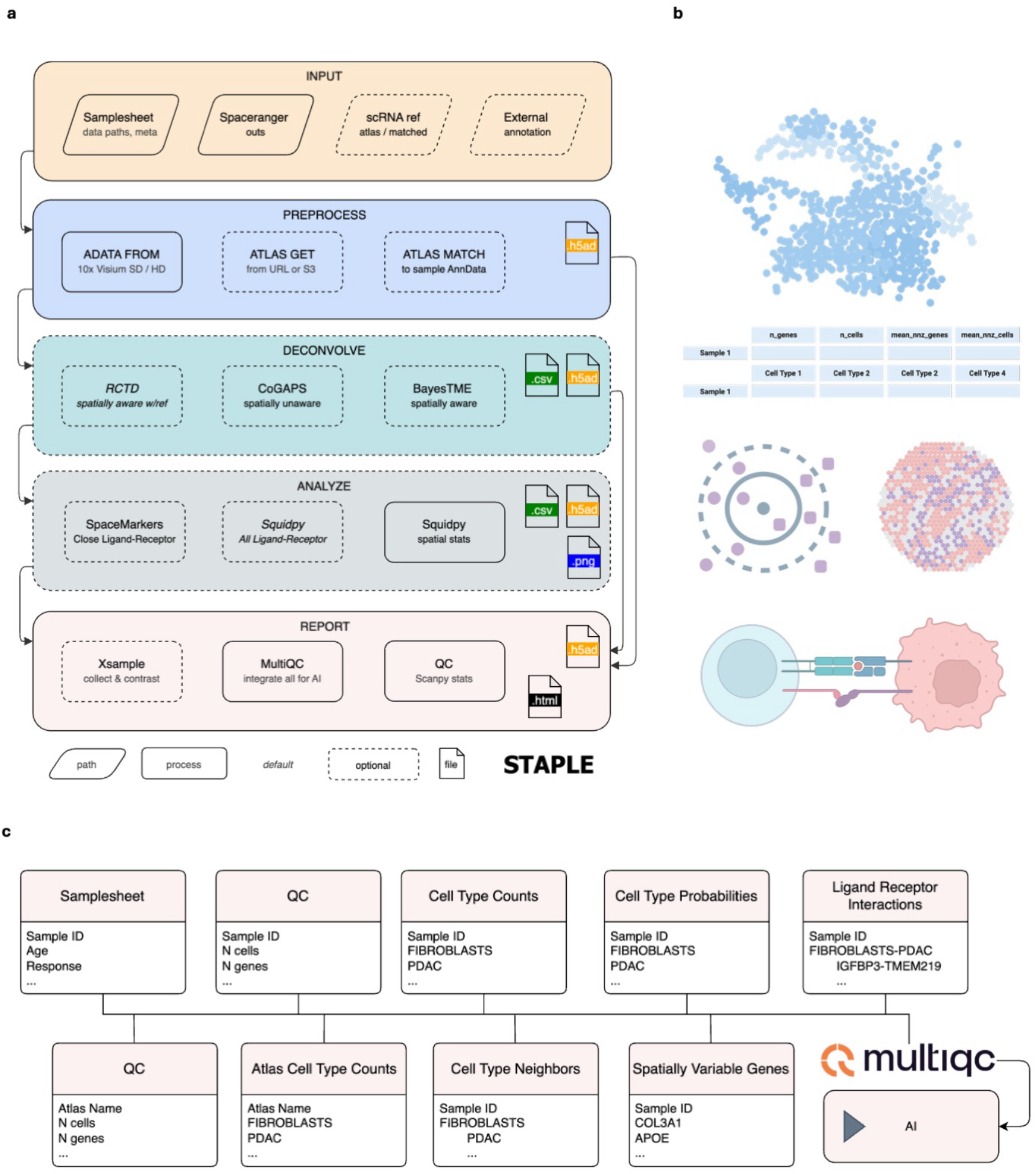
Overview of the pipeline. In STAPLE, containerized tools are orchestrated by Nextflow and use AnnData as the main integration format alongside the tool-agnostic outputs saved for reference. **a**. Pipeline schematics. Inputs to the pipeline are 10X space ranger outputs for Visium SD and Visium HD, optional annotated matched single cell RNAseq data or external barcode-level annotation files, all denoted by rhomboids. Inputs are linked to sample metadata through the sample sheet. In case a single reference atlas is used, it can be provided as a path to the local file system or the cloud (URL or S3, e.g. CellxGene). Data moves sequentially through processes, and processes produce outputs to be consumed by QC/reporting for cross-sample comparison. All metadata present in the sample sheet is carried across the pipeline for downstream correlative analysis. **b**. Graphical representation of outputs from STAPLE. For each sample, cell type annotation (RCTD, BayesTME or CoGAPS), QC (scanpy), cell-type and gene level spatial analysis metrics (Squidpy), ligand-receptor interactions (Squidpy or SpaceMarkers) are computed. **c**. Cross-sample visualization and AI-enabled analysis provided through the MultiQC report. The presence of sample ID in every report table enables AI-assisted multimodal downstream analysis and interpretation.

We demonstrate STAPLE’s broad utility in two biological contexts: a cancer study using Visium high definition (HD) to profile chemotherapy responders and non-responders in pancreatic ductal adenocarcinomas (PDAC)^3^ and in a neuroscience study using Visium standard definition (SD) to characterize nucleus accumbens (NAc)^4^. The PDAC dataset was generated concurrently with STAPLE, therefore the pipeline was optimized for analysis of this clinical cohort and demonstrates the full capacity of STAPLE by automatically annotating cell types, estimating spatial statistics, performing ligand-receptor inference and summarizing results into a comprehensive report that includes comparison between treatment response groups (Figure 2; Supplements 1-4). In this case, we lack a biological ground truth and therefore confirm the findings of these analyses with a large language model (LLM, in this case GPT 5.2 through M365 Copilot) and an expert clinical review. NAc tissue characterization study provides an independent validation of the STAPLE pipeline, in a well-characterized and previously validated dataset. STAPLE analysis of 38 Visium SD human samples generated in the NAc study annotated cell types, computed spatial statistics, and estimated ligand receptor interaction for each sample in under two hours (Supplements 5 and 6). To benchmark functional performance, we applied the LLM and demonstrate that the STAPLE output report is highly concordant with the originally published results (Supplement 7).

**Fig. 2.**
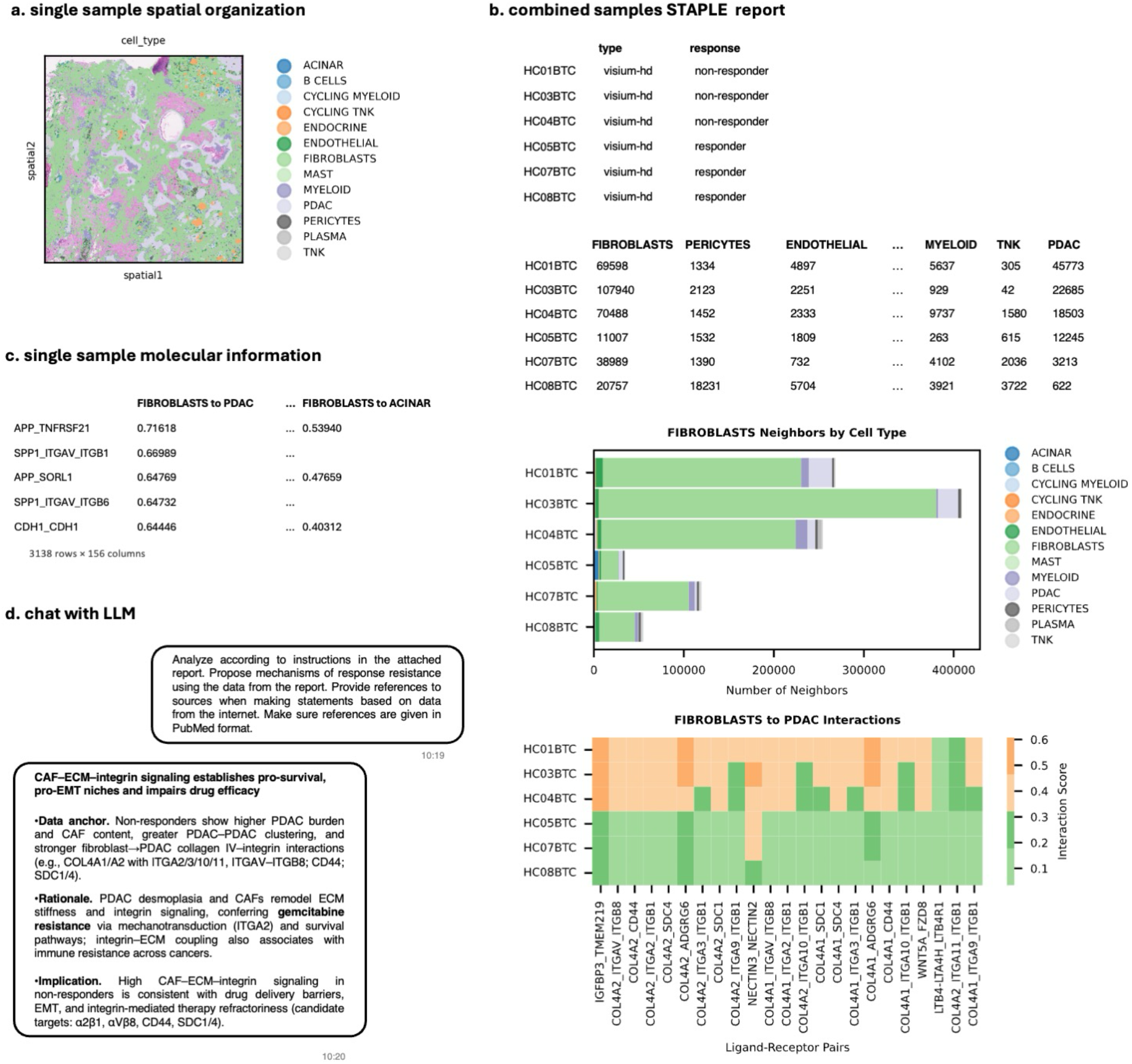
STAPLE applied to the PDAC dataset, selected outputs. Individual modules yield per-sample outputs that contain fragmented characteristics of a data set: **a**. spatial organization, called cell types overlaid on an H&E image shown; **b**. molecular information, illustrated by a fragment of ligand-receptor analysis results These outputs are multi-dimensional and need further treatment to be useful for higher level conclusions. **c**. Sample metadata and sample level outputs are integrated through MultiQC report that provides means for interactive analysis, data visualization, and export to AI models. **d**. Copilot successfully links response status from the sample sheet to other aggregated data tables from the MultiQC report and uses the identified Ligand-Receptor interactions to query the Web for information supporting the findings.

The first step of STAPLE analysis is data ingestion. STAPLE currently accepts spot-based 10X Visium SD, binned Visium HD, and segmented Visium HD data but its modular design and internal data framework make it flexible to additional formats. Following nf-core^5^ convention, STAPLE inputs are read from a sample sheet specifying the sample ID and 10X SpaceRanger^6^ results directory, as well as optional columns for an annotated single-cell RNAseq reference (atlas) and user-provided metadata. Adding metadata columns (such as tissue or organ type, disease status, or treatment regime) instructs STAPLE to preserve those variables in the context of its entire execution and automatically perform contrast analyses on them (Methods). The reference is used for cell typing and deconvolution, and may be specified as a local file, URL or an S3 path in Amazon Cloud Object Storage. This makes it easy to run STAPLE against multiple atlases, even online resources like CellxGene^7^. Online atlases are first downloaded and their cell type labels normalized, before matching against the ST data to ensure consistent gene identifiers are used throughout.

Following ingestion, STAPLE performs cell typing of the ST data in one of 3 ways: (1) label transfer from cell types in the reference atlases, (2) using RCTD; or (3) using one of the reference-free methods (CoGAPS, BayesTME); or a custom annotation provided by the user (Methods). Both of our PDAC (Fig 2 and Supplement 1) and NAc analyses (Supplement 5) use RCTD cell type annotation with custom reference atlases tailored to the biology. Once cell types are annotated, sample characterization and analysis are performed, including the computation of cell type neighborhoods via statistical modules in Squidpy^8^ (Fig 2a); and ligand-receptor interaction performed by Squidpy or SpaceMarkers (Fig. 2b).

Upon completion, results from individual tools are collated into comprehensive reports extending the MultiQC^9^ package from quality control to applications for analytical assessment (shown for PDAC in Fig. 2c). In this case, we leverage MultiQC’s ability to effortlessly combine custom inputs into interactive, AI-ready reports to enable cross-sample analysis in STAPLE. When simply feeding data to MultiQC does lacks interpretability (such as long per-sample lists of spatially variable genes, or complex ligand-receptor interaction objects), STAPLE modules run statistical analyses using sample level metadata (such as case/control or responder/non-responder) and route the contrasted and filtered results back to MultiQC for interactive display.

Individual results within the MultiQC reports are relatively simple flat tables, but deriving a holistic picture from them still poses a time-consuming challenge due to their inherent heterogeneity. This is mitigated by the provenance STAPLE maintains as tools execute along its workflow. Data at each stage of analysis are linked by sample ID, cell type, gene name, and (optionally) matching metadata. This clean, structured data framework enables AI assistants to perform credible first-pass interpretation of the combined report; initiated directly from the built-in MultiQC integration or by export to an LLM of choice (shown for PDAC in Fig. 2d). For full reproducibility, STAPLE outputs all software versions, and produces an execution report summarizing the command(s) used, CPU, RAM and time consumed by each process (Supplements 2 and 6 for PDAC and NAc, respectively).

In summary, we have described STAPLE, a comprehensive, AI-assisted pipeline for systematic spatial transcriptomic analysis; and demonstrated its broad applicability in two distinct research domains, cancer and neuroscience. We discussed how the modular architecture and data framework of STAPLE powers innovative use of an existing reporting tool, MultiQC, to mechanize literature-backed interpretation and first-pass biologically relevant review, by an LLM, directly from automated ST pipeline results. The relative ease with which STAPLE enables AI to be leveraged by non-experts underscores the transformative potential of AI for translational research.

While not compromising rigor, STAPLE promises to reduce fragmentation of the analysis tool chains often used in ST analysis, increase the scalability and repeatability of end-to-end ST workflows, reveal biological signal faster, and sharply decrease analysis turnaround time. The workflow replaces human-intensive manual labor with automated tooling and human-guided machine intelligence to interpret and assess analysis reports.

Future versions of STAPLE will better catalogue samples annotated by reference-free methods, expand its suite of statistical metrics and cross-sample analyses, further streamline reporting for manual and AI review, support more input data formats, and integrate with more LLMs. At the same time, agentic AI methods are emerging to further automate analysis pipelines^10^. The scripting and robust data and metadata handling in pipelines can constrain these AI-derived ST analysis to better ensure reproducibility and grounding in the clinical context.

## Methods

During STAPLE development we contributed code to two open-source reference-free deconvolution tools (CoGAPS^11^, BayesTME^12^) and containerized the reference-based RCTD^13^ deconvolution tool. For ligand-receptor analysis we contributed to SpaceMarkers^14^ and reused tools from Scverse^15^. New Nextflow modules were developed for quality control (QC) summaries, comparing findings across multiple samples, and automatically downloading reference data from CellxGene^7^.

### Inputs and data handling

STAPLE follows the nf-core sample sheet convention for workflow inputs. Sample sheet should contain following columns: *sample, data_directory, expression_profile. sample* is the unique identifier of the sample and is used to link separate tool outputs in the final report. *data_directory* is the path to the SpaceRanger outputs for Visium SD, Visium HD or Visium HD with cell segmentations. *expression_profile* (optional) is a path to sample-level single-cell RNAseq annotated reference. Additional columns may be provided and will be treated as meta data and propagate to the final sample-level outputs and the cross-sample reports.

ST data are ingested using *spatialdata*.*io* readers of scverse^15^, which provide standardized access to multiple spatial omics formats.

AnnData^16^ serves is the central data integration format. Per-sample and per-method AnnData files are saved throughout the workflow, all single method annotations are merged in the final output objects. This design enables independent sample-level processing and delayed aggregation, allowing the pipeline to scale efficiently across large datasets.

### STAPLE outputs

Outputs from STAPLE are described in detail in the accompanying documentation according to the nf-core^5^ template (available from https://github.com/break-through-cancer/staple). Sample-level outputs are stored in folders organized by the names of the tools that generated those, and the sample ID. Raw outputs are provided, as well as selected outputs are aggregated, stored in the final Anndata objects and served in the folder called “staple”.

### Quality control

Sample-level QC summaries for both input spatial data and single-cell atlas data include the number of genes, number of cells (bins) and mean number of genes by counts, mean number of cells by counts and mean total nonzero gene counts using scanpy ^17^.

QC module for cell annotation results adds mean probability of the assigned cell type and counts of cell types per sample, which help evaluate the quality of cell-type assignments and inform downstream analyses.

We also created scripts (available from https://github.com/FertigLab/spatial-test) to subset the publicly available Visium SD and Visium HD datasets and generate test datasets for STAPLE.

### Atlas-derived cell type annotation across samples

RCTD^13^ is selected as the default cell type annotation method in STAPLE based on its high performance across benchmarking studies^18 19^. RCTD^13^ requires reference single-cell RNA-seq gene expression and annotated cell types (atlas). We have containerized RCTD (Dockerfile available from https://github.com/break-through-cancer/btc-containers/tree/main/rctd) and introduced data transformations that allow running RCTD with AnnData input objects as a Nextflow module.

A single atlas can be used to perform cell type annotation across all the samples when specified via STAPLE parameter, or a separate reference can be tied to each ST dataset through the sample sheet, for example when analyzing samples from different tissues, different disease types, or having a matched atlas available. Specific atlases used in our sample pancreatic cancer and brain datasets are provided in our descriptions of these datasets.

When an online location is provided as a source for reference atlas, python packages *boto3* or *requests* are used to automatically download it. STAPLE utility modules are then applied to clean cell the type names and match gene identifiers across the ST and atlas data sets. The gene matching logic computes overlaps between gene names, gene IDs across ST data and the atlas and selects the one identifier where the overlap is larger.

Cell type annotations are assigned to each cell or spot according to the maximum weight in RCTD results.

### Unsupervised cell type annotations in individual samples

STAPLE features two reference-free latent space inference methods, CoGAPS^11^, and BayesTME^12^. Nextflow modules and containers created for both tools upstream in their code repositories, permitting to run them as parts of STAPLE pipeline, or as standalone workflows. Our pipeline is set up to run both methods independently for both methods, providing sample-specific cellular annotations. Annotations are assigned to each spot or cell according to the greatest weight of a feature in the tool’s output. The cellular features learned from any of these methods are further input for within-sample analyses, including spatial neighborhood statistics and ligand–receptor inference.

Our cross-sample analysis relies on shared cellular labels between samples and is thus limited for application in these reference free methods. Nonetheless, the resulting ligand-receptor interaction outputs are incorporated into MultiQC for visualization across samples, but interpretation is limited due to the lack of routine automated analysis pipelines for cross-sample matching of cellular annotations.

### Bring your own cell type annotation

Users may supply pre-computed cell-type probabilities for each spatial barcode obtained externally. In this mode, STAPLE bypasses the cell type annotation step and executes all downstream modules, maintaining the same reporting structure. The required format is a csv file where the first column contains barcodes, and the rest of the columns contain cell proportions estimated for a given barcode by an external method, column names being the cell type names. Examples of this format, generating cell proportions from histologic or spatial proteomics data, are available in the provided references^20^.

### Spatial analysis

We containerized Squidpy^8^ (container image available at https://github.com/break-through-cancer/btc-containers/tree/main/scverse) and implemented it as a module in STAPLE. The Squidpy functions *gr*.*interaction_matrix* and *gr*.*nhood_enrichment* are used to compute per-sample cell type neighborhoods, *gr*.*co_occurrence* to compute pairwise spatial pairwise correlation, *gr*.*spatial_autocorr* for spatially variable genes, *gr*.*centrality_scores* for degree centrality, and *pl*.*spatial_scatter* produces scatter plots. Default parameters are used.

Cell type neighborhood and spatially variable genes are aggregated across samples and displayed in the final MultiQC report. Spatially variable genes output is contrasted using metadata variables before inclusion in the final report.

In addition to Squidpy, we include our spatially aware cell-cell communication method SpaceMarkers^14^ through a Nextflow module and a container contributed to the upstream repository (from https://github.com/DeshpandeLab/SpaceMarkers). This method explicitly estimates transcriptional changes resulting from pairwise spatial overlap between cellular populations. In this implementation, SpaceMarkers uses the overlapping regions as the spatial context for gene expression variation to infer Interaction Marker (IM) scores developed recently to extend the method for both Visium SD and Visium HD data^20^.

### Ligand-receptor interactions

Two ligand-receptor interaction analysis modules are implemented in STAPLE. SpaceMarkers was containerized and a Nextflow module was created upstream as described above, and Squidpy *gr*.*ligrec* method was implemented as a Nextflow module in STAPLE.

In this implementation, SpaceMarkers combines the IM-scores of ligands and the cell-type specificity of receptors to infer ligand-receptor interactions. SpaceMarkers supports an undirected analysis for Visium SD as previously published^14^ and a directed analysis developed recently for Visium HD^3^, owing to near single-cell resolution. Currently, both implementations use spot-based analyses for these inferences.

Squidpy *gr*.*ligrec* uses single cell-type label and identifies interactions between cell-type pairs as described previously^8^. Spatial context must then be derived from spatial neighborhood analysis in this case.

STAPLE harmonizes outputs from both tools to ensure consistent downstream reporting. Full ligand-receptor interaction tables are saved to the final AnnData objects, while top-interacting pairs are filtered and presented in the MultiQC report.

Interpreting ligand–receptor results is challenging due to their high dimensionality; curated databases still yield thousands of potential interactions, and the directional nature of signaling further expands the space.

To manage this complexity, STAPLE applies one of two strategies. When a contrasting metadata variable is available, the workflow uses it to perform a t-test for each ligand– receptor and cell-type pair, corrects for multiple testing, and reports the top interactions that differ across groups. When no such variable exists, STAPLE instead ranks interactions by their mean score across samples and returns the top overall interactions. Then maximum number of interactions to show in the report is regulated by a parameter and defaults to 100.

### Cross-sample comparisons

Individual tool outputs are collected in standardized AnnData object for further bespoke analysis. Sample-level outputs are further aggregated in MultiQC nf-core module through its *custom_data* functionality. Sample sheet is provided as-is after validation and used to give context to the users and AI models. Sample ID interlinks the individual methods results for comprehensive evaluation.

QC metrics for ST inputs and atlases are computed by the QC module per-sample during the pipeline execution. A configuration file describes the logic for incorporating these metrics in the MultiQC report.

The *xsample* (cross-sample) module defines functions for exploring metadata variables that could be used for contrasts given a list of incoming AnnData files, implements statistical testing (t-test using *scipy*.*stats*.*ttest_ind* function), and functions for exporting the ligand-receptor, spatial autocorrelation, and neighborhood analysis to the MultiQC report.

When the number of distinct cell types is large, it may result in underpowered analyses and render cell types of interest hard to find in the reports. STAPLE offers a “spotlight” mode that restricts analysis to user-selected cell types, reducing multiple-testing burden and improving interpretability

### Reporting and AI integration

Upon completion of the sample-level modules, metrics that are created by these modules are collected and combined, together with the sample sheet, into a single interactive MultiQC report that natively supports AI integration. This native AI support includes querying AI models directly from the html MultiQC report by providing API keys to the supported LLMs or manually export the data in a condensed form for use with an LLM of choice. We have used the latter option to and employed the M365 Copilot interface provided by the University of Maryland, Baltimore to ensure that the data in the report are secured with enterprise data protection (see https://learn.microsoft.com/en-gb/copilot/microsoft-365/enterprise-data-protection). The report contains following sections (per sample): sample sheet, input summary, output cell types, cell type probabilities, Moran’s I interactions, ligand-receptor interactions, spatial neighborhood for all cell types called. Each section of the report is a data table accompanied by a name and a short description that provides additional context to AI.

### Application to PDAC dataset

STAPLE was applied to spatial transcriptomics of Visium HD of PDAC dataset using the Cirro cloud platform using the reference RNAseq data from^21^ and preprocessing as described in the original data generation study^3^, producing the sample-level outputs, aggregated MultiQC report (Supplement 1) and the workflow provenance report (Supplement 2).

The MultiQC report was provided to GPT 5.22 via M365 Copilot together with a brief prompt (Fig. 2d). The LLM automatically identified the key elements of the report and performed basic cohort-level contrasts without explicit instruction, generating an initial summary of possible response resistance mechanisms based on the data from the report (Supplement 3).

The summary was reviewed by a pathologist, its general usability confirmed and exploration of the results was proposed. A follow-up prompt yielded expanded analysis containing candidate perturbation agents associated with ligand-receptor pairs identified in the initial results (Supplement 4).

### Application to NAc dataset

For the NAc dataset (38 Visium SD samples), we reformatted GEO files (GSE307586), into 10X Visium Space Ranger format using a custom *write_10x_h5* function, available from https://github.com/FertigLab/staple-paper.

Reference atlas was downloaded from https://released-taxonomies-802451596237-us-west-2.s3.us-west-2.amazonaws.com/HMBA/BasalGanglia/BICAN_05072025_pre-print_release/Human_HMBA_basalganglia_AIT_pre-print.h5ad and subset to retain only NAc region.

Sample sheet was created from GEO files using a custom python script (Code availability). STAPLE was executed using a SLURM cluster at the Institute for Genome Sciences. Workflow output is available in Supplement 5 and provenance report in Supplement 6.

We provided Copilot with the MultiQC report and a prompt requesting comparison between STAPLE-derived findings and the original NAc study. The resulting AI-generated analysis was saved as Supplement 7.

## Supporting information

Supplement 1 - PDAC study STAPLE report

Supplement 2 - PDAC study workflow report

Supplement 3 - PDAC study AI main summary

Supplement 4 - PDAC study AI follow-up summary

Supplement 5 - NAc study STAPLE report

Supplement 6 - NAc study workflow report

Supplement 7 - NAc study AI summary

## Code Availability

STAPLE code and documentation are available from https://github.com/break-through-cancer/staple, the versions used in this publication are available in supplements 1 and 5. Custom codes used to download and preprocess NAc study inputs are available from https://github.com/FertigLab/staple-paper.

## Data Availability

All data used in analysis are available from the original studies^3 4^.

## Acknowledgement and funding

The authors thank Sushma Nagaraj, Ahmed Elhossiny, William Freed-Pastor, Genevieve Stein-O’Brien, Michael F. Ochs and Jesse Boehm for feedback. Funding was provided from Break Through Cancer Data Science and Demystifying Pancreatic Cancer Treatments TeamLabs. In addition, this study was supported by NIH/NCI U24CA284156 (E.J.F., A.D.), U01CA294548 (L.D.W. and E.J.F.), U54CA274371 (E.J.F., L.D.W., M.O.), R37CA271186 (W.T.), U54CA271186 (W.T.), P30CA008748 (MSKCCC), P30CA134274 (UMB), P30CA006973 (JHU), P30CA14051 (Kock Institute, MIT); NIH/NIA U54AG079779 (E.J.F.); NIH / NIDA R01 1DA063090-01 (B.H.); the Lustgarten Foundation (E.J.F., L.T.K., D.S., A.D.); Johns Hopkins University Data Science and Artificial Intelligence institute Demonstration Award (AK); Amazon; the Cancer AI Alliance; the Maurice Campbell Initiative at Memorial Sloan Kettering Cancer Center (WT); and the Maryland Cigarette Restitution Fund (A.D., E.J.F.).

## Data Science Hub TeamLab

Rameen Beroukhim

Leslie Cope

Atul Deshpande

Alexander Favorov

Elana J. Fertig

André Forjaz

Nancy Y. Jung

Rachel Karchin

Luciane T. Kagohara

Ashley L Kiemen

Stuart Levine

Dmitrijs Lvovs

Michael Noble

Wungki Park

Jeffrey Quinn

Sohrab Shah

Dimitrios N. Sidiropoulos

Brandon G Smaglo

Wesley Tansey

Linghua Wang

Meredith Wetzel

Charles A Whittaker

## Demystifying Pancreatic Cancer Therapies TeamLab

Vincent Bernard,

Atul Deshpande,

Elana J. Fertig,

André Forjaz,

William Freed-Pastor,

Luciane T. Kagohara,

Rajya Kappagantula,

Ashley L Kiemen,

Dmitrijs Lvovs,

Anirban Maitra,

Michael Noble,Dimitrios N. Sidiropoulos,Meredith Wetzel,

Laura D Wood

