## Supplement 1 - PDAC study STAPLE report for "STAPLE: automating spatial transcriptomics analysis and AI interpretation"

A modular tool to aggregate results from bioinformatics analyses across many samples into a single report.

This report has been generated by the STAPLE analysis pipeline.

Report generated on 2025-12-21, 21:21 UTC based on data in: `/tmp/nxf.itzmpXxnEn`

Copy report prompt

#### Sample Sheet

The sample sheet provided as input to the pipeline.

Copy table | Configure columns | Scatter plot | Violin plot | Export as CSV... | Showing 6/6 rows and 5/5 columns. | Copy Prompt

| sample | file | type | response | data_directory | expression_profile |
|---|---|---|---|---|---|
| HC01BTC_visiumHD | samplesheet.csv | visium-hd | non-responder | s3://project-5678660b-483c-470c-a0b1-2476fb01fa56/datasets/e4b04b | |
| HC03BTC_visiumHD | samplesheet.csv | visium-hd | non-responder | s3://project-5678660b-483c-470c-a0b1-2476fb01fa56/datasets/2b620c | |
| HC04BTC_visiumHD | samplesheet.csv | visium-hd | non-responder | s3://project-5678660b-483c-470c-a0b1-2476fb01fa56/datasets/1fd526 | |
| HC05BTC_visiumHD | samplesheet.csv | visium-hd | responder | s3://project-5678660b-483c-470c-a0b1-2476fb01fa56/datasets/f88669 | |
| HC07BTC_visiumHD | samplesheet.csv | visium-hd | responder | s3://project-5678660b-483c-470c-a0b1-2476fb01fa56/datasets/7dc585 | |
| HC08BTC_visiumHD | samplesheet.csv | visium-hd | responder | s3://project-5678660b-483c-470c-a0b1-2476fb01fa56/datasets/a3a9b3 | |

#### Atlas Summary

Summary of the cells and genes in the atlas. If same atlas is used for multiple samples, table rows will be repeated.

Copy table | Configure columns | Scatter plot | Violin plot | Export as CSV... | Showing 1/1 rows and 5/5 columns. | Copy Prompt

| Sample | n_genes | n_cells | mean_genes_by_counts | mean_cells_by_counts | mean_total_nnz_counts |
|---|---|---|---|---|---|
| lovelessAtlas_FULLreAnnot_P | 36 601 | 562 321 | 1 300.9 | 19 986.3 | 1.6 |

#### Atlas Cell Types

Summary of the cell types in the atlas.

Copy table | Configure columns | Scatter plot | Violin plot | Export as CSV... | Showing 1/1 rows and 6/13 columns. | Copy Prompt

| Sample | PDAC | TNK | FIBROBLASTS | MYELOID | ENDOTHELIAL | PERICYTES |
|---|---|---|---|---|---|---|
| lovelessAtlas_FULLreAnn | 227 293 | 92 517 | 61 957 | 54 465 | 31 908 | 31 487 |

#### Input Summary

Summary of the AnnData object right after it was read.

Copy table | Configure columns | Scatter plot | Violin plot | Export as CSV... | Showing 6/6 rows and 5/5 columns. | Copy Prompt

| Sample | n_genes | n_cells | mean_genes_by_counts | mean_cells_by_counts | mean_total_nnz_counts |
|---|---|---|---|---|---|
| HC01BTC_visiumHD | 18 085 | 169 550 | 267.6 | 2 508.9 | 1.2 |
| HC03BTC_visiumHD | 18 085 | 175 561 | 176.9 | 1 717.5 | 1.3 |
| HC04BTC_visiumHD | 18 085 | 163 383 | 187.0 | 1 689.7 | 1.2 |
| HC05BTC_visiumHD | 18 085 | 164 263 | 97.4 | 885.0 | 2.4 |
| HC07BTC_visiumHD | 18 085 | 175 561 | 82.6 | 802.3 | 1.2 |
| HC08BTC_visiumHD | 18 085 | 142 989 | 91.7 | 725.0 | 1.2 |

### Output Cell Types

Assigned cell types based on deconvolution / cell typing results.

Copy table | Configure columns | Scatter plot | Violin plot | Export as CSV... | Showing 6/6 rows and 6/13 columns. | Copy Prompt

| Sample | FIBROBLASTS | PERICYTES | ENDOTHELIAL | MYELOID | TNK | PLASMA |
|---|---|---|---|---|---|---|
| HC01BTC_visiumHD | 69 598 | 1 334 | 4 897 | 5 637 | 305 | 409 |
| HC03BTC_visiumHD | 107 940 | 2 123 | 2 251 | 929 | 42 | 219 |
| HC04BTC_visiumHD | 70 488 | 1 452 | 2 333 | 9 737 | 1 580 | 2 070 |
| HC05BTC_visiumHD | 11 007 | 1 532 | 1 809 | 263 | 615 | 41 |
| HC07BTC_visiumHD | 38 989 | 1 390 | 732 | 4 102 | 2 036 | 400 |
| HC08BTC_visiumHD | 20 757 | 18 231 | 5 704 | 3 921 | 3 722 | 1 817 |

### Cell Type Probabilities

Average max cell type probabilities used to assign the cell types (greater is better).

Copy table | Configure columns | Scatter plot | Violin plot | Export as CSV... | Showing 6/6 rows and 6/13 columns. | Copy Prompt

| Sample | FIBROBLASTS | ACINAR | B CELLS | PDAC | CYCLING MYELOID | CYCLING TNK |
|---|---|---|---|---|---|---|
| HC01BTC_visiumHD | 0.6 | 0.5 | 0.3 | 0.7 | 0.4 | 0.3 |
| HC03BTC_visiumHD | 0.7 | 0.4 | 0.5 | 0.6 | 0.4 | |
| HC04BTC_visiumHD | 0.6 | 0.4 | 0.4 | 0.7 | 0.4 | 0.3 |
| HC05BTC_visiumHD | 0.5 | 0.8 | 0.3 | 0.6 | 0.3 | 0.3 |
| HC07BTC_visiumHD | 0.6 | 0.4 | 0.4 | 0.5 | 0.4 | 0.3 |
| HC08BTC_visiumHD | 0.6 | 0.8 | 0.4 | 0.6 | 0.4 | 0.4 |

### Moran I Interactions

Mean interaction across samples shown as no differential interactions were found between ['HC03BTC_visiumHD', 'HC01BTC_visiumHD', 'HC04BTC_visiumHD'] and ['HC05BTC_visiumHD', 'HC08BTC_visiumHD', 'HC07BTC_visiumHD'].

Sorted by sample | Clustered | Copy prompt | Export...

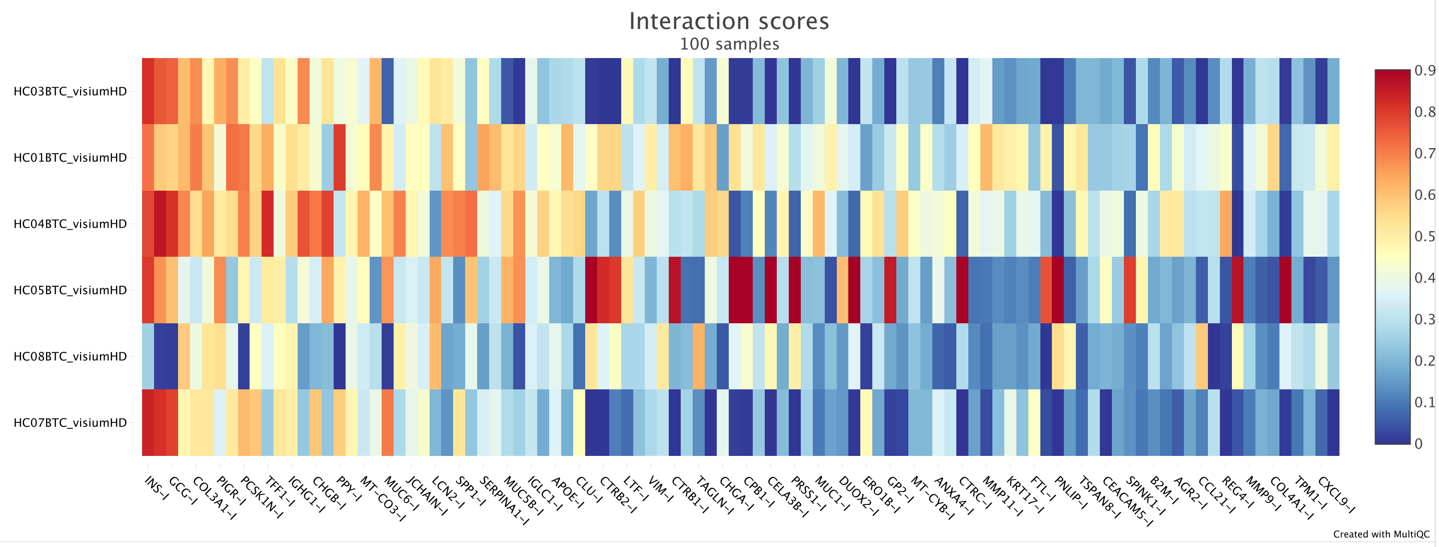

### Spatial Neighbors

Cell type immediate neighborhood across samples.

Percentages | PDAC | FIBROBLASTS | Copy prompt | Export...

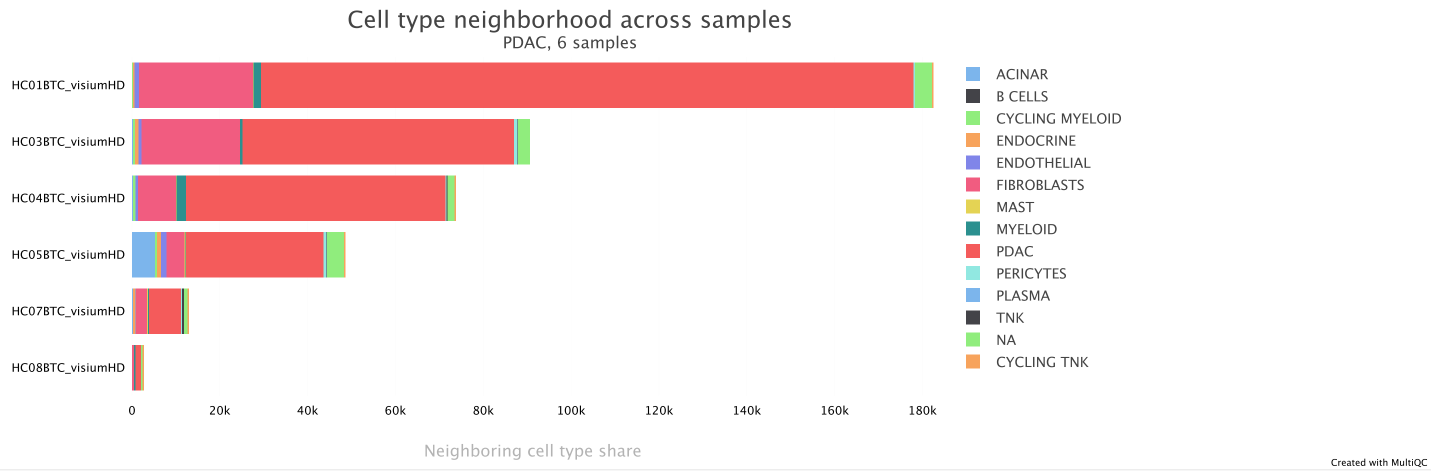


### Spacemarkers Lrscores Interactions

Differential interactions between ['HC03BTC_visiumHD', 'HC01BTC_visiumHD', 'HC04BTC_visiumHD'] and ['HC05BTC_visiumHD', 'HC08BTC_visiumHD', 'HC07BTC_visiumHD']. Top 100 shown sorted by adjusted p-value between 2.34e-02 and 4.81e-02.

Sorted by sample | Clustered | Copy prompt | Export...

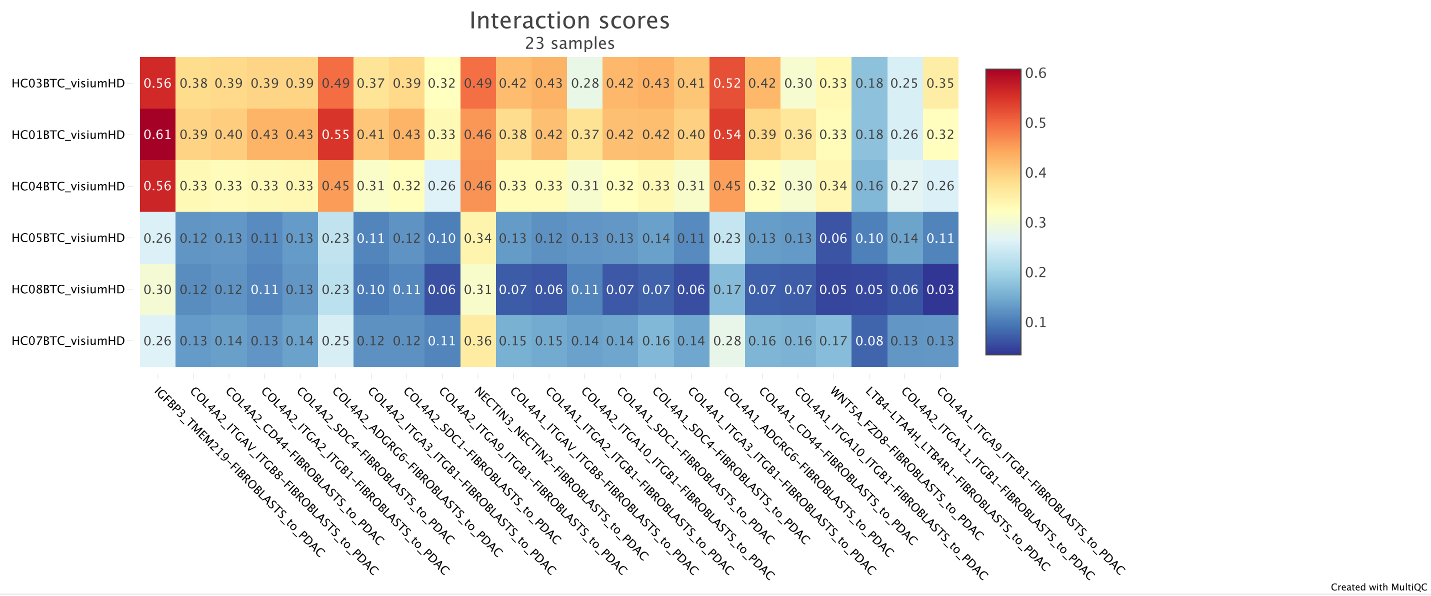

### Software Versions

Software Versions lists versions of software tools extracted from file contents.

Copy table

| Group | Software | Version |
|---|---|---|
| ADATA_ADD_METADATA | anndata | 0.12.6 |
| ADATA_FROM_VISIUM_HD | spatialdata_io | 0.5.1 |
| | squidpy | 1.6.6.dev24+g32789aefd |
| ATLAS_GET | boto3 | 1.41.2 |
| | requests | 2.32.5 |
| ATLAS_MATCH | anndata | 0.12.6 |
| | numpy | 2.3.5 |
| QC | anndata | 0.12.6 |
| | pandas | 2.3.3 |
| RCTD | Matrix | 1.7-4 |
| | R | 4.5.2 |
| | reticulate | 1.44.1 |
| | spacexr | 2.2.1 |
| SQUIDPY_SPATIAL_PLOTS | anndata | 0.12.6 |
| | numpy | 2.3.5 |
| | python | 3.11.14 |
| | squidpy | 1.6.6.dev24+g32789aefd |
| STAPLE_XSAMPLE | anndata | 0.12.6 |
| | json | 2.0.9 |
| | numpy | 2.3.5 |
| | pandas | 2.3.3 |
| | scipy | 1.16.3 |
| Workflow | Nextflow | 25.04.8 |
| | STAPLE | v2.0.0-gb5f784d |
| process | Biobase | 2.70.0 |
| | BiocGenerics | 0.56.0 |
| | CellChat | 2.2.0 |
| | R | 4.5.2 |
| | SpaceMarkers | 2.1.0 |
| | dplyr | 1.1.4 |
| | effsize | 0.8.1 |
| | generics | 0.1.4 |
| | ggplot2 | 4.0.0 |
| | igraph | 2.2.1 |
