## Supplement 2 - PDAC study workflow report for "STAPLE: automating spatial transcriptomics analysis and AI interpretation"

### Nextflow workflow report

#### [cirro-e5597-f3b76] *(resumed run)*

Workflow execution completed successfully!

**Run times**

21-Dec-2025 21:14:38 - 21-Dec-2025 21:21:40 (duration: **7m 2s**)

3 s... 87 cached

**Nextflow command**

```
nextflow run break-through-cancer/btc-spatial-pipelines/main.nf -r 116-downstream-cross-sample-analysis -params-file params.json -resume
49861b5c-d8ba-4baf-9a06-20cf5ddf9664 -name cirro-e5597-f3b76 -latest -ansi-log false -with-report
/opt/work/834c4cf3814122d7a431217bf9244dd2/artifacts/report.html -with-timeline
/opt/work/834c4cf3814122d7a431217bf9244dd2/artifacts/timeline.html -with-dag
/opt/work/834c4cf3814122d7a431217bf9244dd2/artifacts/flowchart.svg -config nextflow-override.config
```

| | |
|---|---|
| **CPU-Hours** | 324.3 (100% cached) |
| **Launch directory** | /opt/work/834c4cf3814122d7a431217bf9244dd2 |
| **Work directory** | s3://project-5678660b-483c-470c-a0b1-2476fb01fa56-scratch/workdir |
| **Project directory** | /root/.nextflow/assets/break-through-cancer/btc-spatial-pipelines |
| **Script name** | main.nf |
| **Script ID** | 5a95f7d52dc1cd094f2b1163a4ea12a7 |
| **Workflow session** | 49861b5c-d8ba-4baf-9a06-20cf5ddf9664 |
| **Workflow repository** | https://github.com/break-through-cancer/btc-spatial-pipelines , revision 116-downstream-cross-sample-analysis (commit hash b5f784d3bcd7a24320d6814efd475fb68efcee71 ) |
| **Workflow profile** | standard |
| **Nextflow version** | version 25.04.8, build 5956 (06-10-2025 21:19 UTC) |

#### Resource Usage

These plots give an overview of the distribution of resource usage for each process.

##### CPU

Raw Usage | % Allocated

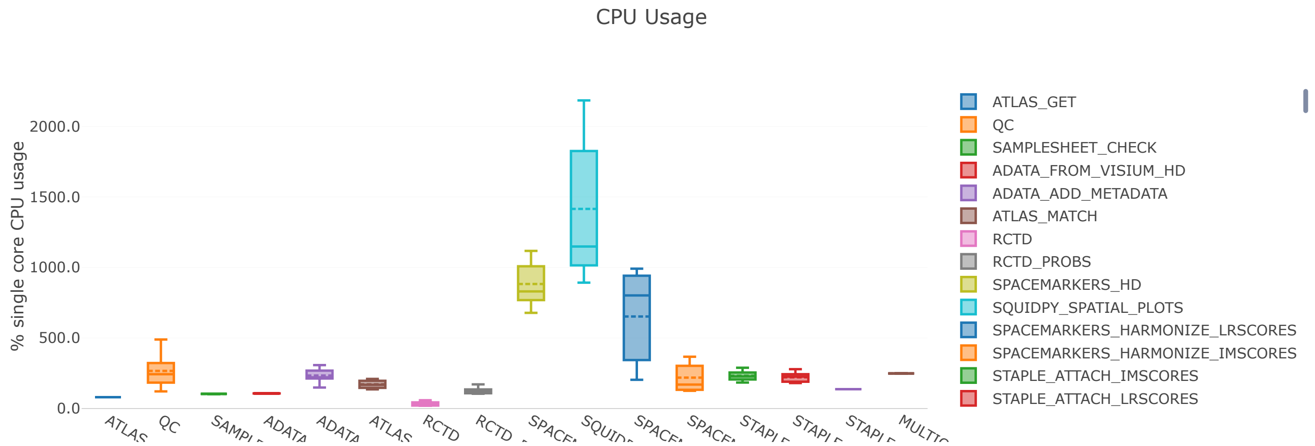

#### Memory

Physical (RAM) | Virtual (RAM + Disk swap) | % RAM Allocated

Physical Memory Usage

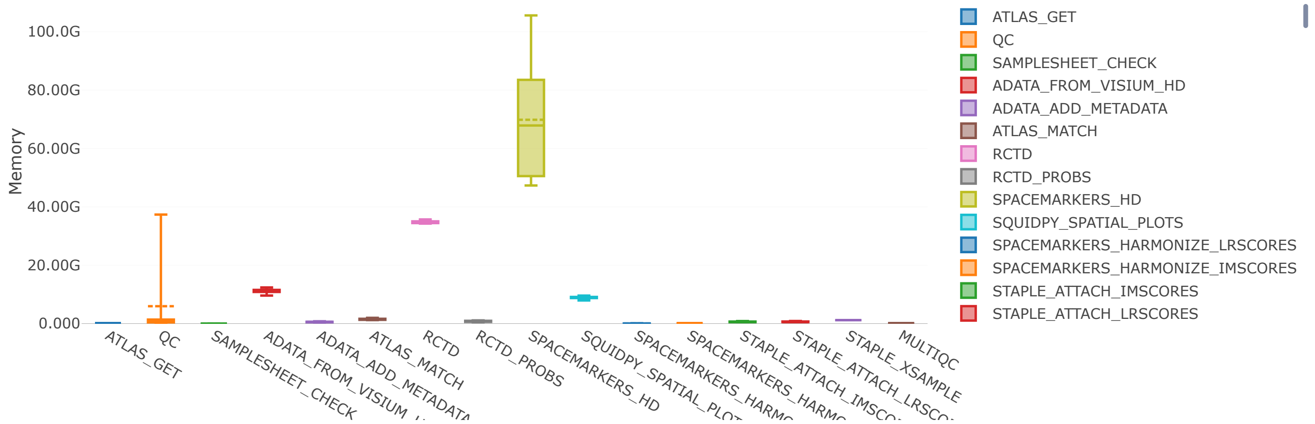


#### Job Duration

Raw Usage | % Allocated

Task execution real-time

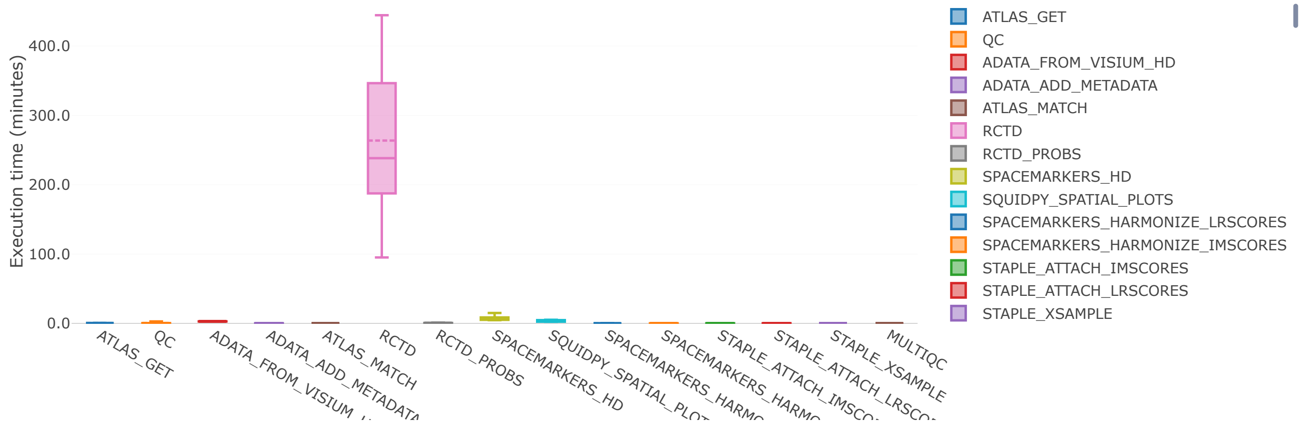


## I/O

Read | Write

Number of bytes read

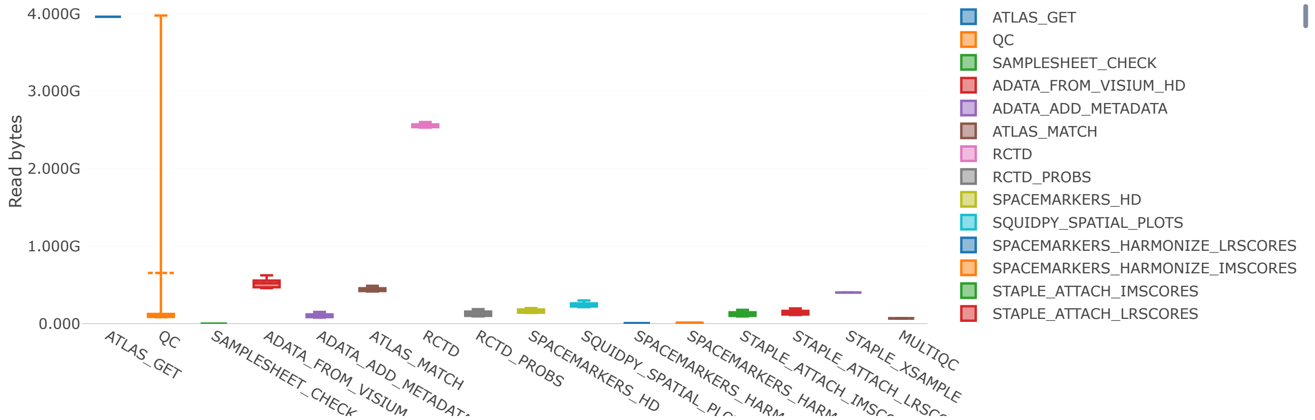

### Tasks

This table shows information about each task in the workflow. Use the search box on the right to filter rows for specific values. Clicking headers will sort the table by that value and scrolling side to side will reveal more columns.

Values shown as: Human readable

Show 25 entries Filter: Metrics Metadata All Search:

| task_id | process | tag | status | hash | allocated cpus | %cpu | allocated memory | %mem |
|---|---|---|---|---|---|---|---|---|
| 1 | BTC:STAPLE:LOAD_DATASET:ATLA | - | CACHED | f0/ad8a96 | 2 | 80.4 | 12.000 GB | 0.7 |
| 2 | BTC:STAPLE:INPUT_CHECK:SAMPL | cirro-samplesheet.csv | COMPLETED | 9f/750c77 | 1 | 104.0 | 8.000 GB | 0.0 |
| 3 | BTC:STAPLE:QC | - | CACHED | 8f/d05d70 | 4 | 221.5 | 32.000 GB | 0.8 |
| 4 | BTC:STAPLE:QC | - | CACHED | d2/fd29a1 | 8 | 121.7 | 64.000 GB | 47.2 |
| 5 | BTC:STAPLE:LOAD_DATASET:ADAT | - | CACHED | 6d/c09173 | 4 | 107.1 | 32.000 GB | 8.8 |
| 6 | BTC:STAPLE:LOAD_DATASET:ADAT | - | CACHED | 30/cb4f74 | 4 | 107.0 | 32.000 GB | 8.1 |
| 7 | BTC:STAPLE:LOAD_DATASET:ADAT | - | CACHED | 3a/cdc6d7 | 4 | 107.4 | 32.000 GB | 7.7 |
| 8 | BTC:STAPLE:LOAD_DATASET:ADAT | HC03BTC_visiumHD | CACHED | 6a/146b55 | 4 | 221.7 | 32.000 GB | 0.4 |
| 9 | BTC:STAPLE:QC | - | CACHED | 8c/0de8de | 4 | 246.6 | 32.000 GB | 1.0 |
| 10 | BTC:STAPLE:LOAD_DATASET:ADAT | - | CACHED | ee/6bbf8d | 4 | 107.1 | 32.000 GB | 7.7 |
| 11 | BTC:STAPLE:LOAD_DATASET:ADAT | HC01BTC_visiumHD | CACHED | 84/be887b | 4 | 149.0 | 32.000 GB | 1.1 |
| 12 | BTC:STAPLE:QC | - | CACHED | b7/04306b | 4 | 240.3 | 32.000 GB | 0.8 |

| | | | | | | | | |
|---|---|---|---|---|---|---|---|---|
| 14 | BTC:STAPLE:QC | - | CACHED | 35/bfb2dc | 4 | 257.2 | 32.000 GB | 1.1 |
| 15 | BTC:STAPLE:LOAD_DATASET:ATLA | HC01BTC_visiumHD | CACHED | 03/84ccac | 4 | 136.0 | 32.000 GB | 2.8 |
| 16 | BTC:STAPLE:LOAD_DATASET:ADAT | HC04BTC_visiumHD | CACHED | cd/24d895 | 4 | 216.9 | 32.000 GB | 0.4 |
| 17 | BTC:STAPLE:LOAD_DATASET:ADAT | - | CACHED | 24/ae9d20 | 4 | 108.1 | 32.000 GB | 6.6 |
| 18 | BTC:STAPLE:LOAD_DATASET:ADAT | HC05BTC_visiumHD | CACHED | db/1e054f | 4 | 284.6 | 32.000 GB | 0.3 |
| 19 | BTC:STAPLE:LOAD_DATASET:ATLA | HC03BTC_visiumHD | CACHED | 69/3ef862 | 4 | 178.1 | 32.000 GB | 1.2 |
| 20 | BTC:STAPLE:QC | - | CACHED | 12/d303fb | 4 | 268.1 | 32.000 GB | 1.0 |
| 21 | BTC:STAPLE:DECONVOLVE:RCTD | HC01BTC_visiumHD | CACHED | c4/279a54 | 12 | 58.9 | 128.000 GB | 14.3 |
| 22 | BTC:STAPLE:LOAD_DATASET:ADAT | HC07BTC_visiumHD | CACHED | 7d/b56ed7 | 4 | 211.7 | 32.000 GB | 0.6 |
| 23 | BTC:STAPLE:QC | - | CACHED | 39/6e52d3 | 4 | 262.3 | 32.000 GB | 0.5 |
| 24 | BTC:STAPLE:DECONVOLVE:RCTD | HC03BTC_visiumHD | CACHED | 80/aa3158 | 12 | 30.1 | 128.000 GB | 15.7 |
| 25 | BTC:STAPLE:LOAD_DATASET:ATLA | HC04BTC_visiumHD | CACHED | 48/b93d35 | 4 | 142.6 | 32.000 GB | 2.4 |

Showing 1 to 25 of 90 entries

Previous 1 2 3 4 Next
