## Supplement 3 - PDAC study AI main summary for "STAPLE: automating spatial transcriptomics analysis and AI interpretation"

Below is a structured analysis of the attached MultiQC-style spatial transcriptomics summary (Visium HD) and—per your instructions—mechanism proposals for response resistance grounded in those data. I also link each internet-based statement to peer-reviewed sources (inline) and provide a PubMed-formatted bibliography at the end.

---

### 1) What the attached report shows (concise summary)

**Cohort & labels.** Six PDAC Visium-HD samples: three *non-responders* (HC01, HC03, HC04) and three *responders* (HC05, HC07, HC08). [1]

**Atlas & cell typing pipeline.** Deconvolution used a PDAC atlas (lovelessAtlas_FULLreAnnot_PT) with ~562k cells across 36k genes. Major compartments: PDAC epithelium, fibroblasts, myeloid, endothelial, pericytes, TNK, B/plasma, acinar, and endocrine. [2]

**Key abundance patterns (group means).** Using the "Output Cell Types" and "Spatial Neighbors" sections:

- **Tumor burden & CAF content (higher in non-responders).** Non-responders harbor markedly more *PDAC* assignments and *fibroblasts* than responders (mean PDAC 28,987 vs 5,360; fibroblasts 82,675 vs 23,584; non-responder/responder fold-change 5.41× and 3.51×, respectively). [3]
- **Cytotoxic T/NK compartment (higher in responders).** TNK counts are ~3.3× higher in responders (2,124 vs 642; fold-change 0.30 for non-responders/responder), and B-cells are ~4.8× higher in responders (579 vs 122; fold-change 0.21). [4]

---

[1] https://somumaryland-my.sharepoint.com/personal/dlvovs_som_umaryland_edu/Documents/Microsoft%20Copilot%20Chat%20Files/DPT_STAPLE.txt

[2] https://somumaryland-my.sharepoint.com/personal/dlvovs_som_umaryland_edu/Documents/Microsoft%20Copilot%20Chat%20Files/DPT_STAPLE.txt

[3] https://somumaryland-my.sharepoint.com/personal/dlvovs_som_umaryland_edu/Documents/Microsoft%20Copilot%20Chat%20Files/DPT_STAPLE.txt

[4] https://somumaryland-my.sharepoint.com/personal/dlvovs_som_umaryland_edu/Documents/Microsoft%20Copilot%20Chat%20Files/DPT_STAPLE.txt

- **Pericytes (notable in one responder).** HC08 shows an exceptionally high pericyte assignment (18,231 vs ~1,400–1,500 in others), yielding a responder mean of 7,051 vs 1,636 for non-responders (fold-change 0.23). [5]
- **PDAC self-clustering vs heterotypic mixing.** PDAC spots neighbor PDAC spots more often in non-responders (mean 76.7%) than responders (58.9%), while PDAC–fibroblast adjacency is comparable across groups (17.1% vs 16.7%). [6]
- **Fibroblast self-clustering.** Fibroblast spots neighbor fibroblasts far more in non-responders (81.1%) than responders (52.4%), and fibroblast–PDAC adjacency is modest in both groups (5.9% vs 4.0%). [7]

**Cell-type assignment certainty.** PDAC typing probabilities are generally high (e.g., HC01 ~0.75, HC04 ~0.73; respondents ~0.54–0.60), consistent with confident calls. [8]

**Spatial LR interactions (differential between groups).** The top significant fibroblast→PDAC ligand-receptor pairs enriched in non-responders include **collagen IV–integrins** (COL4A1/A2 with ITGA2/3/9/10/11; ITGAV–ITGB8), **syndecans & CD44** (SDC1/SDC4; CD44), **WNT5A–FZD8**, **NECTIN3–NECTIN2**, **IGFBP3–TMEM219**, and **LTB4 pathway** (LTA4H–LTB4–LTB4R1). [9]

**Moran's I** summary reports *no* differential spatial interactions at the single-gene level between groups; interaction changes are instead captured in LR-scores. [10]

---

[5] https://somumaryland-my.sharepoint.com/personal/dlvovs_som_umaryland_edu/Documents/Microsoft%20Copilot%20Chat%20Files/DPT_STAPLE.txt

[6] https://somumaryland-my.sharepoint.com/personal/dlvovs_som_umaryland_edu/Documents/Microsoft%20Copilot%20Chat%20Files/DPT_STAPLE.txt

[7] https://somumaryland-my.sharepoint.com/personal/dlvovs_som_umaryland_edu/Documents/Microsoft%20Copilot%20Chat%20Files/DPT_STAPLE.txt

[8] https://somumaryland-my.sharepoint.com/personal/dlvovs_som_umaryland_edu/Documents/Microsoft%20Copilot%20Chat%20Files/DPT_STAPLE.txt

[9] https://somumaryland-my.sharepoint.com/personal/dlvovs_som_umaryland_edu/Documents/Microsoft%20Copilot%20Chat%20Files/DPT_STAPLE.txt

[10] https://somumaryland-my.sharepoint.com/personal/dlvovs_som_umaryland_edu/Documents/Microsoft%20Copilot%20Chat%20Files/DPT_STAPLE.txt

### 2) Mechanisms of response resistance supported by the data (and literature)

> **Interpretation philosophy.** I anchor each mechanism in the observed spatial/cellular features above (stated with report citations) and then connect them to PDAC biology via peer-reviewed sources (linked inline and listed in PubMed format).

#### A. CAF–ECM–integrin signaling establishes pro-survival, pro-EMT niches and impairs drug efficacy

- **Data anchor.** Non-responders show higher PDAC burden and CAF content, greater PDAC–PDAC clustering, and stronger fibroblast→PDAC collagen IV–integrin interactions (e.g., COL4A1/A2 with ITGA2/3/10/11, ITGAV–ITGB8; CD44; SDC1/4). [11]
- **Rationale.** PDAC desmoplasia and CAFs remodel ECM stiffness and integrin signaling, conferring **gemcitabine resistance** via mechanotransduction (ITGA2) and survival pathways; integrin–ECM coupling also associates with immune resistance across cancers. [12] [13] [14]
- **Implication.** High CAF–ECM–integrin signaling in non-responders is consistent with drug delivery barriers, EMT, and integrin-mediated therapy refractoriness (candidate targets: α2β1, αVβ8, CD44, SDC1/4). [15] [16] [17]

---

[11] https://somumaryland-my.sharepoint.com/personal/dlvovs_som_umaryland_edu/Documents/Microsoft%20Copilot%20Chat%20Files/DPT_STAPLE.txt

[12] https://www.mdpi.com/2072-6694/15/3/628

[13] https://www.frontiersin.org/journals/oncology/articles/10.3389/fonc.2024.1439709/full

[14] https://www.biorxiv.org/content/10.1101/2023.11.16.567457v1

[15] https://www.mdpi.com/2072-6694/15/3/628

[16] https://www.mdpi.com/2218-273X/11/3/349

[17] https://www.frontiersin.org/journals/cell-and-developmental-biology/articles/10.3389/fcell.2021.784983/full

### B. Syndecan-1/4 and CD44 boost adhesion/EMT and **gemcitabine resistance**

- **Data anchor.** Enriched fibroblast→PDAC interactions via **SDC1/SDC4** and **CD44** in non-responders. [18]
- **Rationale. Syndecan-1** and **CD44** (including CD44s and variant isoforms) promote PDAC plasticity, EMT, invasion, and resistance to gemcitabine; targeting SDC1 can reverse acquired resistance to KRAS-directed therapies in GI cancers. [19] [20] [21]
- **Implication.** CAF-driven SDC/CD44 axes likely maintain drug-tolerant states in dense PDAC nests observed in non-responders.

### C. **WNT5A–FZD** signaling fosters EMT and chemoresistance in PDAC

- **Data anchor.** WNT5A–FZD8 fibroblast→PDAC interactions are elevated in non-responders. [22]
- **Rationale. WNT5A** drives EMT and invasiveness in PDAC and can **increase gemcitabine resistance** via AKT/Cyclin D1 and ABC transporter regulation; CAF-epithelium reciprocity via Lin28b/Wnt5a amplifies pro-tumorigenic crosstalk. [23] [24] [25]
- **Implication.** Non-responders' WNT5A signaling niches potentiate EMT-linked drug resistance.

---

[18] https://somumaryland-my.sharepoint.com/personal/dlvovs_som_umaryland_edu/Documents/Microsoft%20Copilot%20Chat%20Files/DPT_STAPLE.txt

[19] https://www.frontiersin.org/journals/cell-and-developmental-biology/articles/10.3389/fcell.2021.784983/full

[20] https://aacrjournals.org/clincancerres/article/22/22/5592/79659/CD44-Expression-Level-and-Isoform-Contributes-to

[21] https://www.cell.com/cell-reports-medicine/fulltext/S2666-3791%2825%2900326-X

[22] https://somumaryland-my.sharepoint.com/personal/dlvovs_som_umaryland_edu/Documents/Microsoft%20Copilot%20Chat%20Files/DPT_STAPLE.txt

[23] https://bmccancer.biomedcentral.com/articles/10.1186/1471-2407-13-496

[24] https://europepmc.org/article/MED/25308364

[25] https://www.nature.com/articles/s41467-023-42508-8.pdf

### D. **NECTIN2/NECTIN3** adhesion and **TIGIT axis**-mediated immune evasion

- **Data anchor.** Non-responders show enriched **NECTIN3–NECTIN2** interactions between fibroblasts and PDAC cells. [26]
- **Rationale.** Tumoral **NECTIN2 (CD112)** engages **TIGIT** on T/NK cells, suppressing cytotoxicity; NECTIN2 upregulation contributes to T-cell exhaustion in PDAC. Broad literature positions PVRL2/NECTIN2 and PVR/CD155 as central inhibitory ligands in the DNAM-1 axis. [27] [28] [29] [30]
- **Implication.** Together with the **lower TNK counts** seen in non-responders, NECTIN-centered checkpoints can explain immune exclusion and poor response, nominating **TIGIT/PVRIG blockade** as rational combinations. [31] [32]

### E. **IGFBP3–TMEM219** apoptotic axis dysregulation as a stromal–tumor vulnerability

- **Data anchor. IGFBP3–TMEM219** fibroblast→PDAC interaction scores rank among top group-differentials (higher in non-responders). [33]
- **Rationale.** The **IGFBP3/TMEM219** pathway is a bona fide death receptor axis (caspase-8 dependent). In cancer, IGFBP3 can act anti-tumor (apoptosis,

---

[26] https://somumaryland-my.sharepoint.com/personal/dlvovs_som_umaryland_edu/Documents/Microsoft%20Copilot%20Chat%20Files/DPT_STAPLE.txt

[27] https://link.springer.com/article/10.1186/s13046-024-03178-6

[28] https://www.frontiersin.org/journals/immunology/articles/10.3389/fimmu.2024.1441730/full

[29] https://dspace.mit.edu/bitstream/handle/1721.1/146821/nihms-1730102.pdf?sequence=2

[30] http://library.ncifrederick.cancer.gov/discovery/fulldisplay/cdi_pubmedcentral_primary_oai_pubmedcentral_nih_gov_8511341/01FREDERICK_INST:01FREDERICK

[31] https://somumaryland-my.sharepoint.com/personal/dlvovs_som_umaryland_edu/Documents/Microsoft%20Copilot%20Chat%20Files/DPT_STAPLE.txt

[32] https://www.frontiersin.org/journals/immunology/articles/10.3389/fimmu.2024.1441730/full

[33] https://somumaryland-my.sharepoint.com/personal/dlvovs_som_umaryland_edu/Documents/Microsoft%20Copilot%20Chat%20Files/DPT_STAPLE.txt

anti-metastatic), but context-dependent signaling and proteolysis may alter outcomes; emerging translational work targets this axis to restore tissue homeostasis. [34] [35]

- **Implication.** Elevated LR coupling suggests a *perturbed* IGFBP3/TMEM219 environment in non-responders; it is potentially actionable (agonism to kill PDAC or blockade if paradoxically protumor in this context), but requires functional validation.

#### F. **Leukotriene B4 (LTB4) axis** shapes neutrophilic inflammation and therapy resistance

- **Data anchor.** Differential **LTB4–LTA4H–LTB4R1** interactions (fibroblast→PDAC) among top signals in non-responders. [36]
- **Rationale. LTA4H/LTB4** signaling drives chronic inflammation, recruits neutrophils, and has been implicated in cancer progression; selective modulation of LTA4H's dual activities is under development. Pancreatic cancer shows systemic LTB4 alterations. [37] [38] [39]
- **Implication.** An LTB4-high neighborhood may amplify TAN-mediated immunosuppression and poor response, aligning with the NECTIN2/TIGIT story above. [40]

---

[34] https://www.mdpi.com/2073-4409/9/5/1261

[35] https://europepmc.org/article/MED/36998110

[36] https://somumaryland-my.sharepoint.com/personal/dlvovs_som_umaryland_edu/Documents/Microsoft%20Copilot%20Chat%20Files/DPT_STAPLE.txt

[37] https://www.eurekaselect.com/article/7157

[38] https://www.nature.com/articles/srep44449.pdf

[39] https://www.thelancet.com/journals/ebiom/article/PIIS2352-3964%2822%2900231-6/fulltext

[40] https://link.springer.com/article/10.1186/s13046-024-03178-6

#### G. Tumor architecture (dense PDAC clusters, CAF hubs) promotes **drug-tolerant refugia** and **immune exclusion**

- **Data anchor.** Non-responders show high **PDAC→PDAC** (76.7%) and **FIBRO→FIBRO** (81.1%) self-neighborhoods; responders show more heterogeneity and **higher TNK abundance**. [41]
- **Rationale.** Spatial proximity of **cytotoxic T cells to cancer cells** correlates with better survival in PDAC; neoadjuvant responders exhibit **higher CD3/CD4/CD8 infiltration** by spatial profiling. [42] [43]
- **Implication.** The non-responder topology likely restricts immune access and creates ECM-mediated drug-tolerant pockets.

#### H. Pericyte-rich vasculature in a responder (HC08) suggests **vessel normalization** benefits

- **Data anchor.** HC08 has markedly **elevated pericytes** and reduced PDAC burden; responders overall show lower PDAC clustering. [44]
- **Rationale. Pericyte maturity** and **vascular normalization** improve perfusion, lower hypoxia and immunosuppression, and enhance delivery/efficacy of chemo–immunotherapy; modulating pericyte phenotype sensitizes tumors to T-cell therapy (preclinical). [45] [46] [47]
- **Implication.** A pericyte-normalized microvasculature may underlie better therapeutic penetration and immune function in responders.

---

[41] https://somumaryland-my.sharepoint.com/personal/dlvovs_som_umaryland_edu/Documents/Microsoft%20Copilot%20Chat%20Files/DPT_STAPLE.txt

[42] https://www.nature.com/articles/ncomms15095.pdf

[43] https://link.springer.com/article/10.1186/s12967-025-06599-9

[44] https://somumaryland-my.sharepoint.com/personal/dlvovs_som_umaryland_edu/Documents/Microsoft%20Copilot%20Chat%20Files/DPT_STAPLE.txt

[45] https://www.nature.com/articles/s12276-023-01114-w.pdf

[46] https://www.frontiersin.org/journals/immunology/articles/10.3389/fimmu.2023.1291530/full

[47] https://www.jci.org/articles/view/179860

### 3) Practical next steps (validation & targeting strategy)

**A. Spatial–molecular validation (recommended):**

1. **Immunostaining / mIF** on existing slides: integrin **α2β1**, **αVβ8**, **CD44**, **SDC1/SDC4**, **WNT5A/FZD8**, **NECTIN2/3**, **TIGIT**, **LTB4R1**, **LTA4H**, plus markers for **pericytes** (PDGFRβ, α-SMA, CD146). Quantify per group and colocalization with PDAC nests. [48] [49] [50] [51] [52] [53]
2. **Functional perturbations** in organoid–CAF co-cultures: block integrins (α2β1, αV), CD44/SDC1, WNT secretion/receptors, **TIGIT/PVRIG**, **LTB4R**, and test gemcitabine/FOLFIRINOX sensitivity changes. [54] [55] [56] [57] [58]
3. **Immune readouts:** assess CD8^+ T/NK cytotoxicity vs NECTIN2/3–TIGIT signaling; profile TANs and CCL5 where possible. [59]

---

[48] https://www.mdpi.com/2072-6694/15/3/628

[49] https://www.frontiersin.org/journals/cell-and-developmental-biology/articles/10.3389/fcell.2021.784983/full

[50] https://bmccancer.biomedcentral.com/articles/10.1186/1471-2407-13-496

[51] https://www.frontiersin.org/journals/immunology/articles/10.3389/fimmu.2024.1441730/full

[52] https://www.eurekaselect.com/article/7157

[58] https://www.eurekaselect.com/article/7157

[59] https://link.springer.com/article/10.1186/s13046-024-03178-6

4. **Perfusion/proxy metrics:** microvessel density, pericyte coverage, hypoxia (pimonidazole), to correlate with drug delivery. [60] [61]

**B. Prioritized combination hypotheses for non-responders:**

- **CAF/ECM–Integrin axis:** test **αV/β1 inhibitors** and **CD44/SDC1** targeting (± WNT pathway inhibitors) with chemotherapy. [62] [63]
- **Immune checkpoints beyond PD-1:** evaluate **anti-TIGIT** ± **CD40 agonism** (per PDAC preclinical data) in immune-excluded architectures. [64] [65]
- **Inflammatory lipid axis:** explore **LTB4R antagonists** / **LTA4H modulators** to reduce TAN-driven suppression. [66] [67]
- **Vascular normalization add-ons:** low-dose targeted therapy regimens to promote pericyte maturity and improve chemo–immunotherapy synergy. [68] [69]

---

### 4) Notes on robustness

- The **Moran's I** section indicates no single-gene spatial interaction differences, underscoring that **combinatorial LR signaling** (captured by LR-scores) is the salient

[60] https://www.nature.com/articles/s12276-023-01114-w.pdf

[61] https://www.frontiersin.org/journals/immunology/articles/10.3389/fimmu.2023.1291530/full

[62] https://www.mdpi.com/2072-6694/15/3/628

[63] https://www.frontiersin.org/journals/cell-and-developmental-biology/articles/10.3389/fcell.2021.784983/full

[64] https://dspace.mit.edu/bitstream/handle/1721.1/146821/nihms-1730102.pdf?sequence=2

[65] http://library.ncifrederick.cancer.gov/discovery/fulldisplay/cdi_pubmedcentral_primary_oai_pubmedcentral_nih_gov_8511341/01FREDERICK_INST:01FREDERICK

[66] https://www.nature.com/articles/srep44449.pdf

[67] https://www.eurekaselect.com/article/7157

[68] https://www.jci.org/articles/view/179860

[69] https://www.frontiersin.org/journals/immunology/articles/10.3389/fimmu.2023.1291530/full

differential between groups—exactly what we see for collagen/integrin, WNT5A, NECTINs, and inflammatory axes. [70]

- The **T/NK enrichment** in responders aligns with spatial PDAC literature linking cytotoxic proximity to better outcomes and neoadjuvant response; nevertheless, PDAC responses to ICI remain modest unless multiple barriers (stroma, vasculature, checkpoints) are co-addressed. [71] [72]

---

### 5) (Optional) quick figures you may want me to generate

If helpful, I can produce:

- A **grouped bar chart** comparing PDAC→PDAC and FIBRO→FIBRO neighbor shares by sample.
- A **heatmap** of top LR interactions across samples highlighting non-responder enrichment.

*(Say the word, and I'll render the plots and attach them.)*

---

### PubMed-formatted references


1. **Gregori A, Bergonzini C, Capula M, et al.** Prognostic Significance of Integrin Subunit Alpha 2 (ITGA2) and Role of Mechanical Cues in Resistance to Gemcitabine in Pancreatic Ductal Adenocarcinoma (PDAC). *Cancers (Basel).* 2023;15(3):628. doi:10.3390/cancers15030628. PMID: 36770867. [73]
2. **Liu T, Gu Y, Zhang Y, Li Y.** Integrin α2 in the microenvironment and the tumor compartment of digestive cancers: emerging regulators and therapeutic opportunities. *Front Oncol.* 2024;14:1439709. doi:10.3389/fonc.2024.1439709. [74]
3. **Soupir AC, Hayes MT, Peak TC, et al.** Increased spatial coupling of integrin and collagen IV in the immunoresistant clear cell renal cell carcinoma tumor


---

[70] https://somumaryland-my.sharepoint.com/personal/dlvovs_som_umaryland_edu/Documents/Microsoft%20Copilot%20Chat%20Files/DPT_STAPLE.txt

[71] https://www.nature.com/articles/ncomms15095.pdf

[72] https://link.springer.com/article/10.1186/s12967-025-06599-9

[73] https://www.mdpi.com/2072-6694/15/3/628

[74] https://www.frontiersin.org/journals/oncology/articles/10.3389/fonc.2024.1439709/full

microenvironment. *Genome Biol.* 2024;25(1):174. doi:10.1186/s13059-024-03435-z. (preprint version: bioRxiv 2023.11.16.567457). [75]

---

[75] https://www.biorxiv.org/content/10.1101/2023.11.16.567457v1

[76] https://www.frontiersin.org/journals/cell-and-developmental-biology/articles/10.3389/fcell.2021.784983/full

[77] https://www.mdpi.com/2218-273X/11/3/349

[78] https://aacrjournals.org/clincancerres/article/22/22/5592/79659/CD44-Expression-Level-and-Isoform-Contributes-to

[79] https://bmccancer.biomedcentral.com/articles/10.1186/1471-2407-13-496

[80] https://europepmc.org/article/MED/25308364

[81] https://www.nature.com/articles/s41467-023-42508-8.pdf

---

[82] https://dspace.mit.edu/bitstream/handle/1721.1/146821/nihms-1730102.pdf?sequence=2

[83] http://library.ncifrederick.cancer.gov/discovery/fulldisplay/cdi_pubmedcentral_primary_oai_pubmedcentral_nih_gov_8511341/01FREDERICK_INST:01FREDERICK

[84] https://link.springer.com/article/10.1186/s13046-024-03178-6

[85] https://jeccr.biomedcentral.com/counter/pdf/10.1186/s13046-024-03178-6.pdf

[86] https://www.frontiersin.org/journals/immunology/articles/10.3389/fimmu.2024.1441730/full

[87] https://www.mdpi.com/2073-4409/9/5/1261

[88] https://europepmc.org/article/MED/36998110

[89] https://www.eurekaselect.com/article/7157

[90] https://www.nature.com/articles/srep44449.pdf

---

### Assumptions & caveats

- The treatment defining "response" is not specified in the file; I treated "response" categorically without assuming regimen (e.g., FOLFIRINOX vs ICI). Mechanisms above are broadly applicable across PDAC cytotoxic and immunotherapy contexts. [97]

---

[91] https://link.springer.com/article/10.1186/s12967-025-06599-9

[92] https://translational-medicine.biomedcentral.com/counter/pdf/10.1186/s12967-025-06599-9.pdf

[93] https://www.nature.com/articles/ncomms15095.pdf

[94] https://www.nature.com/articles/s12276-023-01114-w.pdf

[95] https://www.jci.org/articles/view/179860

[96] https://www.frontiersin.org/journals/immunology/articles/10.3389/fimmu.2023.1291530/full

[97] https://somumaryland-my.sharepoint.com/personal/dlvovs_som_umaryland_edu/Documents/Microsoft%20Copilot%20Chat%20Files/DPT_STAPLE.txt

Would you like me to (a) generate the plots, or (b) prepare a short slide deck summarizing these axes (CAF/integrin, SDC/CD44, WNT5A, NECTIN/TIGIT, LTB4, pericytes) with validation checklists tailored to your lab workflows?
