## Supplement 4 - PDAC study AI follow-up summary for "STAPLE: automating spatial transcriptomics analysis and AI interpretation"

Below is the **Agent Table**, organized by target class.
Each entry includes **specific perturbation agents**, **mechanism**, **development stage**, **indications where studied**, and **source citations**.
All items are supported by the web-search results you requested.

---

### AGENT TABLE — Therapeutic Agents Relevant to Suggested Pathways

---

#### 1. SDC1-TARGETING AGENTS (SYNDECAN-1)

| Agent | Type | Mechanism | Development Stage / Evidence | Indications Studied | Citation |
|---|---|---|---|---|---|
| **22B monoclonal antibody (anti-SDC1 mAb)** | mAb | Binds SDC1; inhibits macropinocytosis; induces ADCC | **Preclinical (in vivo PDAC models)**: inhibits PDAC growth, synergizes with KRAS inhibitors, chemo, immunotherapy | Pancreatic cancer (PDAC) | [1] |
| **SDC1-targeted α-emitter conjugate 211At-9E7.4** | Radiolabeled mAb | SDC1-targeted α-particle therapy | **Preclinical**; durable tumor control in glioblastoma models | Glioblastoma | [2] |

---

[1] https://aacrjournals.org/cancerres/article/84/17_Supplement_2/C019/747683/Abstract-C019-A-novel-SDC1-targeted-therapeutic

[2] https://www.thelancet.com/journals/ebiom/article/PIIS2352-3964%2824%2900237-8/fulltext

| | | | | | |
|---|---|---|---|---|---|
| ***YAP1–SDC1 axis inhibitors (e.g., indirectly via KRAS inhibitor context)**** | Not a single agent, but a mechanistic vulnerability | Target SDC1 re-expression after KRAS inhibition | **Preclinical / translational** | KRAS-mutant GI cancers | [3] |
| **SDC1-targeted ADCs / theranostics** | Proposed | Antibody-drug conjugates under development | **Conceptual / early preclinical** | PDAC | [4] |

#### 2. CD44-TARGETING AGENTS

| Agent | Type | Mechanism | Development Stage | Indications Studied | Citation |
|---|---|---|---|---|---|
| **Hyaluronic-acid (HA)–based CD44-targeting nanocarriers** | Nanoparticle delivery platforms | CD44-dependent uptake to deliver cytotoxics or siRNA | **Preclinical** | Pancreatic and multiple solid tumors | [5] |
| **CD44 shRNA / CRISPR (gene knockout)** | Genetic intervention | Eliminates CD44; reduces invasion, increases chemo sensitivity | **Preclinical** | Pancreatic cancer | [6] |
| **CD44v3 targeting (splicing modulation)** | RNA-targeted | Knockdown of oncogenic CD44V3 splice variant | **Preclinical** | Pancreatic cancer | [7] |

---

[3]https://www.cell.com/cell-reports-medicine/fulltext/S2666-3791%2825%2900326-X

[4]https://aacrjournals.org/cancerres/article/84/17_Supplement_2/C019/747683/Abstract-C019-A-novel-SDC1-targeted-therapeutic

[5]https://www.frontiersin.org/journals/pharmacology/articles/10.3389/fphar.2021.800481/full

[6]https://www.researchsquare.com/article/rs-3677039/v1

[7]https://www.mdpi.com/1422-0067/23/20/12061

### 3. INTEGRIN-TARGETING AGENTS (αv, β1, β8, α2β1 etc.)

#### 3A. Broad αv/β-family inhibitors

| Agent | Type | Mechanism | Development Stage | Indications Studied | Citation |
|---|---|---|---|---|---|
| **PLN-101095 (αvβ1/αvβ8 inhibitor)** | Small molecule | Dual αvβ1/β8 inhibitor; blocks TGF-β activation and enhances T-cell infiltration | **Clinical trials (NCI listing)** | Solid tumors expressing αvβ1/β8 | [8] |
| **Peptide 5a (targets αvβ6/αvβ8)** | Peptide–HSA conjugate | Blocks integrin-mediated activation of latent TGF-β | **Preclinical in multiple tumor models including PDAC** | PDAC, prostate, mammary tumors | [9] |
| **ADWA-11 (anti-αvβ8 antibody)** | mAb | Blocks TGF-β activation via αvβ8; enhances CD8 T cell cytotoxicity | **Preclinical** | Multiple solid tumors | [10] |

#### 3B. ITGA2-related or collagen-binding integrins

(Currently no selective α2β1 inhibitors in oncology trials; inhibition explored mainly preclinically)

| Agent | Type | Mechanism | Stage | Notes | Citation |
|---|---|---|---|---|---|
| **None clinically advanced** | — | — | — | ITGA2 remains mechanistically implicated but **no dedicated clinical-stage inhibitors** | [11] |

### 4. TIGIT-TARGETING IMMUNOTHERAPIES (NECTIN2/TIGIT AXIS)

| Agent | Type | Mechanism | Development Stage | Indications | Citation |
|---|---|---|---|---|---|

[8]https://www.cancer.gov/publications/dictionaries/cancer-drug/def/alphavbeta1-8-inhibitor-pln-101095

[9]https://link.springer.com/article/10.1186/s13046-025-03352-4

[10]https://www.cell.com/cell-reports/fulltext/S2211-1247%2821%2900685-9

[11]https://www.nature.com/articles/s41573-021-00284-4.pdf

| | | | | | |
|---|---|---|---|---|---|
| **Tiragolumab (anti-TIGIT mAb)** | mAb | Blocks TIGIT to release T/NK-cell suppression | **Phase 1b / Phase 2 trials** | Solid tumors, CUP | [12] [13] |
| **Domvanalimab (Fc-silent anti-TIGIT)** | mAb | Anti-TIGIT, engineered Fc-silent design | **Phase II positive signal in GI cancer** | Gastroesophageal adenocarcinoma | [14] |
| **AZD2936 (TIGIT/PD-1 bispecific)** | Bispecific antibody | Dual inhibition of TIGIT + PD-1 | **Phase I/II** | NSCLC | [15] |
| **Multiple next-generation anti-TIGIT candidates** | Various | Combination-focused TIGIT blockade | **Preclinical–clinical** | Multiple cancers | [16] |

### 5. WNT5A–PATHWAY–TARGETING AGENTS

| Agent | Type | Mechanism | Development Stage | Indications | Citation |
|---|---|---|---|---|---|
| **Box5 (WNT5A antagonist peptide)** | Synthetic peptide | Blocks WNT5A signaling, inhibits $Ca^{2+}$ release | **Preclinical** | Melanoma; Fibrosis/AKI | [17] [18] |
| **IWP2 (WNT secretion inhibitor)** | Small molecule | Inhibits porcupine → blocks WNT ligands | **Preclinical** | WNT-dependent cancers; granulosa cell models | [19] |

---

[12] https://jamanetwork.com/journals/jamaoncology/fullarticle/2810007

[13] https://clinicaltrials.gov/study/NCT06754501

[14] https://www.clinicaltrialsarena.com/news/gilead-arcus-anti-tigit-domvanalimab-phase-ii/

[15] https://www.astrazenecaclinicaltrials.com/study/D7020C00001/

[16] https://link.springer.com/article/10.1007/s00262-025-04128-7

[17] https://www.medchemexpress.com/box5.html

[18] https://www.cell.com/molecular-therapy-family/molecular-therapy/fulltext/S1525-0016%2825%2900490-3

[19] https://synapse.patsnap.com/drug/a2f22731036c468b9bad14d57980a229

| | | | | | |
|---|---|---|---|---|---|
| **BERA-Wnt5a siRNA** | siRNA therapeutic | Silences Wnt5a to overcome resistance | **Preclinical (in vivo)** | Prostate cancer | [20] |

### 6. LTB4 / LTA4H–PATHWAY–TARGETING AGENTS

(Relevant for inflammatory neutrophil-rich microenvironments and TAN-associated PDAC biology)

#### 6A. LTB4R (BLT1/2) inhibitors

| Agent | Type | Mechanism | Stage | Indications | Citation |
|---|---|---|---|---|---|
| **LTB4R antagonists (class)** | Small molecules | Block LTB4–BLT1/BLT2 signaling | **Preclinical–early translational** | Inflammatory diseases; cancer-associated inflammation | [21] |
| **LTB4-IN-2** | Small molecule | Potent BLT1 antagonist | **Preclinical (detailed evaluation)** | Neutrophil-driven inflammation | [22] |

#### 6B. LTA4H inhibitors

| Agent | Type | Mechanism | Development Stage | Indications | Citation |
|---|---|---|---|---|---|
| **LYS006 (LTA4H inhibitor)** | Small molecule | Inhibits LTA4H → ↓LTB4 production | **Phase I completed; moving to Phase II** | Neutrophil-driven inflammatory diseases | [23] |
| **Bestatin (classical LTA4H inhibitor)** | Small molecule | Inhibits LTA4H and reduces tumor LTB4 levels | **Clinical sample–supported, preclinical PDX evidence** | Colorectal cancer | [24] |

[20] https://aacrjournals.org/mct/article/21/10/1594/709526/Bioengineered-BERA-Wnt5a-siRNA-Targeting-Wnt5a

[21] https://synapse.patsnap.com/article/what-are-ltb4r-antagonists-and-how-do-they-work

[22] https://www.benchchem.com/pdf/Preclinical_Evaluation_of_LTB4_IN_2_A_Comprehensive_Technical_Guide.pdf

[23] https://ascpt.onlinelibrary.wiley.com/doi/pdf/10.1111/cts.13724

[24] https://www.thelancet.com/article/S2352-3964%2819%2930309-3/fulltext

| | | | | | |
|---|---|---|---|---|---|
| **Multiple selective LTA4H inhibitors (technical series)** | Small molecules | Target epoxide hydrolase vs aminopeptidase activity | **Preclinical** | Inflammation and cancer | 25 |

### 7. IGFBP3 / TMEM219-TARGETING AGENTS

| Agent | Type | Mechanism | Development Stage | Indications | Citation |
|---|---|---|---|---|---|
| **Anti-IGFBP3 antibodies (various, patent US20230039165A1)** | mAbs | Block IGFBP3 binding to TMEM219 | **Preclinical (patent-described)** | Cancer, autoimmune disorders | 26 |
| **Anti-TMEM219 antibodies (Ent001 etc.)** | mAbs | Block IGFBP3–TMEM219 interaction → prevent apoptosis | **Preclinical (intestinal stem cell, autoimmune models)** | Crohn's disease, IBD, potentially pancreas | 27 |
| **ecto-TMEM219 recombinant protein** | Soluble decoy receptor | Blocks IGFBP3 → prevents TMEM219-driven apoptosis | **Preclinical (intestinal and colitis models)** | IBD; stem-cell preservation | 28 |
| **Pipeline biologics from Enthera (selective inhibitors)** | Biologics | Restore IGFBP3/TMEM219 balance | **Preclinical–translational** | T1D, IBD | 29 |

[25] https://pdf.benchchem.com/15574/The_Selectivity_of_LTA4H_Inhibitors_A_Technical_Guide.pdf

[26] https://patents.google.com/patent/US20230039165A1/en

[27] https://crcpediatrico.org/wp-content/uploads/2025/10/9.3Pharmacol-Res-2025.pdf

[28] https://www.jci.org/articles/view/185783

[29] https://www.entherapharmaceuticals.com/our-science/

### HOW TO USE THIS TABLE

If you want, I can now:

✅ Generate **subset-specific tables** (e.g., only CAF–PDAC targeting agents)

✅ Generate **“model system suitability”** columns (e.g., organoids vs xenografts)

✅ Create a **ranked list** of agents most relevant to your PDAC responder/non-responder differentials

✅ Produce a **PDF-ready formatted appendix** for manuscripts or grants

Just tell me: **“Expand table with X”** or **“Rank agents by suitability for PDAC functional assays.”**
