## Supplement 5 - NAc study STAPLE report for "STAPLE: automating spatial transcriptomics analysis and AI interpretation"

multiqc

A modular tool to aggregate results from bioinformatics analyses across many samples into a single report.

This report has been generated by the STAPLE analysis pipeline.

Report generated on 2026-02-06, 18:29 EST based on data in: `/usr/local/scratch/dlvovs/work/98/29c02d487b3e7e5e960758935d9500`

Copy report prompt

### Sample Sheet

The sample sheet provided as input to the pipeline.

Copy table | Configure columns | Scatter plot | Violin plot | Export as CSV... | Showing 38/38 rows and 7/7 columns. | Copy Prompt

| sample | data_directory | expression_profile | brain | tissue | region | subject_status | age |
|---|---|---|---|---|---|---|---|
| **GSM9227428** | data/GSE307586/samples/GSM9227428 | | Br2720 | nucleus accumbens | A1 | neurotypical control postm[illegible]onor | 48.2 |
| **GSM9227429** | data/GSE307586/samples/GSM9227429 | | Br2720 | nucleus accumbens | B1 | neurotypical control postm[illegible]onor | 48.2 |
| **GSM9227430** | data/GSE307586/samples/GSM9227430 | | Br2720 | nucleus accumbens | C1 | neurotypical control postm[illegible]onor | 48.2 |
| **GSM9227431** | data/GSE307586/samples/GSM9227431 | | Br2720 | nucleus accumbens | D1 | neurotypical control postm[illegible]onor | 48.2 |
| **GSM9227432** | data/GSE307586/samples/GSM9227432 | | Br8492 | nucleus accumbens | A1 | neurotypical control postm[illegible]or | 53.4 |
| **GSM9227433** | data/GSE307586/samples/GSM9227433 | | Br8492 | nucleus accumbens | B1 | neurotypical control postm[illegible]or | 53.4 |
| **GSM9227434** | data/GSE307586/samples/GSM9227434 | | Br8492 | nucleus accumbens | C1 | neurotypical control postm[illegible]or | 53.4 |
| **GSM9227435** | data/GSE307586/samples/GSM9227435 | | Br8492 | nucleus accumbens | D1 | neurotypical control postm[illegible]or | 53.4 |
| **GSM9227436** | data/GSE307586/samples/GSM9227436 | | Br6522 | nucleus accumbens | A1 | neurotypical control postm[illegible]rain donor | 33.4 |
| **GSM9227437** | data/GSE307586/samples/GSM9227437 | | Br6522 | nucleus accumbens | B1 | neurotypical control postm[illegible]rain donor | 33.4 |
| **GSM9227438** | data/GSE307586/samples/GSM9227438 | | Br6522 | nucleus accumbens | C1 | neurotypical control postm[illegible]rain donor | 33.4 |
| **GSM9227439** | data/GSE307586/samples/GSM9227439 | | Br6522 | nucleus accumbens | D1 | neurotypical control postm[illegible]rain donor | 33.4 |
| **GSM9227440** | data/GSE307586/samples/GSM9227440 | | Br6423 | nucleus accumbens | A1 | neurotypical control postm[illegible]nor | 51.7 |
| **GSM9227441** | data/GSE307586/samples/GSM9227441 | | Br6423 | nucleus accumbens | B1 | neurotypical control postm[illegible]nor | 51.7 |
| **GSM9227442** | data/GSE307586/samples/GSM9227442 | | Br8325 | nucleus accumbens | C1 | neurotypical control postm[illegible] | 57.6 |
| **GSM9227443** | data/GSE307586/samples/GSM9227443 | | Br6423 | nucleus accumbens | D1 | neurotypical control postm[illegible]nor | 51.7 |
| **GSM9227444** | data/GSE307586/samples/GSM9227444 | | Br8325 | nucleus accumbens | A1 | neurotypical control postm[illegible] | 57.6 |
| **GSM9227445** | data/GSE307586/samples/GSM9227445 | | Br8325 | nucleus accumbens | B1 | neurotypical control postm[illegible] | 57.6 |
| **GSM9227446** | data/GSE307586/samples/GSM9227446 | | Br8325 | nucleus accumbens | C1 | neurotypical control postm[illegible] | 57.6 |
| **GSM9227447** | data/GSE307586/samples/GSM9227447 | | Br8325 | nucleus accumbens | D1 | neurotypical control postm[illegible] | 57.6 |
| **GSM9227448** | data/GSE307586/samples/GSM9227448 | | Br6432 | nucleus accumbens | A1 | neurotypical control postm[illegible]onor | 48.9 |
| **GSM9227449** | data/GSE307586/samples/GSM9227449 | | Br6432 | nucleus accumbens | B1 | neurotypical control postm[illegible]onor | 48.9 |
| **GSM9227450** | data/GSE307586/samples/GSM9227450 | | Br6432 | nucleus accumbens | C1 | neurotypical control postm[illegible]onor | 48.9 |
| **GSM9227451** | data/GSE307586/samples/GSM9227451 | | Br6432 | nucleus accumbens | D1 | neurotypical control postm[illegible]onor | 48.9 |
| **GSM9227452** | data/GSE307586/samples/GSM9227452 | | Br6471 | nucleus accumbens | A1 | neurotypical control postm[illegible]r | 55.5 |
| **GSM9227453** | data/GSE307586/samples/GSM9227453 | | Br6471 | nucleus accumbens | B1 | neurotypical control postm[illegible]r | 55.5 |
| **GSM9227454** | data/GSE307586/samples/GSM9227454 | | Br6471 | nucleus accumbens | C1 | neurotypical control postm[illegible]r | 55.5 |
| **GSM9227455** | data/GSE307586/samples/GSM9227455 | | Br6471 | nucleus accumbens | D1 | neurotypical control postm[illegible]r | 55.5 |
| **GSM9227456** | data/GSE307586/samples/GSM9227456 | | Br2743 | nucleus accumbens | A1 | neurotypical control postm[illegible] | 61.5 |
| **GSM9227457** | data/GSE307586/samples/GSM9227457 | | Br2743 | nucleus accumbens | B1 | neurotypical control postm[illegible] | 61.5 |
| **GSM9227458** | data/GSE307586/samples/GSM9227458 | | Br2743 | nucleus accumbens | C1 | neurotypical control postm[illegible] | 61.5 |
| **GSM9227459** | data/GSE307586/samples/GSM9227459 | | Br2743 | nucleus accumbens | D1 | neurotypical control postm[illegible] | 61.5 |
| **GSM9227460** | data/GSE307586/samples/GSM9227460 | | Br3942 | nucleus accumbens | A1 | neurotypical control postm[illegible]onor | 47.5 |
| **GSM9227461** | data/GSE307586/samples/GSM9227461 | | Br3942 | nucleus accumbens | B1 | neurotypical control postm[illegible]onor | 47.5 |
| **GSM9227462** | data/GSE307586/samples/GSM9227462 | | Br3942 | nucleus accumbens | C1 | neurotypical control postm[illegible]onor | 47.5 |
| **GSM9227463** | data/GSE307586/samples/GSM9227463 | | Br3942 | nucleus accumbens | D1 | neurotypical control postm[illegible]onor | 47.5 |
| **GSM9227464** | data/GSE307586/samples/GSM9227464 | | Br8667 | nucleus accumbens | B1 | neurotypical control postm[illegible]ain donor | 37.3 |
| **GSM9227465** | data/GSE307586/samples/GSM9227465 | | Br8667 | nucleus accumbens | C1 | neurotypical control postm[illegible]ain donor | 37.3 |

### Atlas Summary

Summary of the cells and genes in the atlas. If same atlas is used for multiple samples, table rows will be repeated.

Copy table | Configure columns | Scatter plot | Violin plot | Export as CSV... | Showing 1/1 rows and 5/5 columns. | Copy Prompt

| Sample | n_genes | n_cells | mean_genes_by_counts | mean_cells_by_counts | mean_total_nnz_counts |
|---|---|---|---|---|---|
| Human_HMBA_basalganglia_A print_NAC | 36 601 | 116 884 | 5 239.4 | 16 731.9 | 4.5 |

### Atlas Cell Types

Summary of the cell types in the atlas.

Copy table | Configure columns | Scatter plot | Violin plot | Export as CSV... | Showing 1/1 rows and 6/33 columns. | Copy Prompt

| Sample | STR D1 MSN | STR D2 MSN | Astrocyte | Oligodendrocyte | STR Hybrid MSN | CN ST18 GABA |
|---|---|---|---|---|---|---|
| Human_HMBA_basalgang print_NAC | 32 743 | 32 219 | 14 136 | 9 901 | 8 528 | 5 338 |

### Input Summary

Summary of the AnnData object right after it was read.

Copy table | Configure columns | Scatter plot | Violin plot | Export as CSV... | Showing 38/38 rows and 5/5 columns. | Copy Prompt

| Sample | n_genes | n_cells | mean_genes_by_counts | mean_cells_by_counts | mean_total_nnz_counts |
|---|---|---|---|---|---|
| GSM9227428 | 36 601 | 4 992 | 1 659.9 | 226.4 | 2.0 |
| GSM9227429 | 36 601 | 4 992 | 2 652.0 | 361.7 | 2.3 |
| GSM9227430 | 36 601 | 4 992 | 2 590.1 | 353.3 | 2.2 |
| GSM9227431 | 36 601 | 4 992 | 2 616.5 | 356.9 | 2.2 |
| GSM9227432 | 36 601 | 4 991 | 1 583.6 | 215.9 | 2.0 |
| GSM9227433 | 36 601 | 4 992 | 1 481.7 | 202.1 | 2.0 |
| GSM9227434 | 36 601 | 4 992 | 1 555.5 | 212.2 | 2.1 |
| GSM9227435 | 36 601 | 4 992 | 1 671.5 | 228.0 | 2.0 |
| GSM9227436 | 36 601 | 4 992 | 2 654.3 | 362.0 | 2.6 |
| GSM9227437 | 36 601 | 4 992 | 2 615.9 | 356.8 | 2.5 |
| GSM9227438 | 36 601 | 4 992 | 2 710.2 | 369.6 | 2.3 |
| GSM9227439 | 36 601 | 4 992 | 2 262.6 | 308.6 | 2.3 |
| GSM9227440 | 36 601 | 4 992 | 1 682.1 | 229.4 | 1.8 |
| GSM9227441 | 36 601 | 4 992 | 1 094.1 | 149.2 | 1.8 |
| GSM9227442 | 36 601 | 4 992 | 2 569.1 | 350.4 | 2.1 |
| GSM9227443 | 36 601 | 4 992 | 2 152.4 | 293.6 | 2.0 |
| GSM9227444 | 36 601 | 4 992 | 2 393.1 | 326.4 | 2.2 |
| GSM9227445 | 36 601 | 4 992 | 2 271.3 | 309.8 | 2.1 |
| GSM9227446 | 36 601 | 4 992 | 2 171.0 | 296.1 | 2.1 |
| GSM9227447 | 36 601 | 4 992 | 2 409.4 | 328.6 | 2.1 |
| GSM9227448 | 36 601 | 4 992 | 2 125.8 | 289.9 | 2.1 |
| GSM9227449 | 36 601 | 4 992 | 1 405.6 | 191.7 | 1.9 |
| GSM9227450 | 36 601 | 4 992 | 875.2 | 119.4 | 1.6 |
| GSM9227451 | 36 601 | 4 992 | 1 825.0 | 248.9 | 2.0 |
| GSM9227452 | 36 601 | 4 992 | 2 023.6 | 276.0 | 1.9 |
| GSM9227453 | 36 601 | 4 992 | 2 160.2 | 294.6 | 1.8 |
| GSM9227454 | 36 601 | 4 992 | 1 634.5 | 222.9 | 1.7 |
| GSM9227455 | 36 601 | 4 992 | 1 797.2 | 245.1 | 1.8 |
| GSM9227456 | 36 601 | 4 992 | 1 688.3 | 230.3 | 2.0 |
| GSM9227457 | 36 601 | 4 992 | 1 463.8 | 199.6 | 1.9 |
| GSM9227458 | 36 601 | 4 992 | 1 067.2 | 145.5 | 2.0 |
| GSM9227459 | 36 601 | 4 992 | 949.5 | 129.5 | 1.9 |
| GSM9227460 | 36 601 | 4 992 | 2 131.7 | 290.7 | 2.2 |
| GSM9227461 | 36 601 | 4 992 | 1 699.6 | 231.8 | 2.2 |
| GSM9227462 | 36 601 | 4 992 | 1 586.8 | 216.4 | 2.1 |
| GSM9227463 | 36 601 | 4 992 | 1 829.1 | 249.5 | 2.1 |
| GSM9227464 | 36 601 | 4 974 | 2 628.0 | 357.1 | 2.5 |
| GSM9227465 | 36 601 | 4 976 | 2 037.3 | 277.0 | 2.4 |

### Output Cell Types

Assigned cell types based on deconvolution / cell typing results.

Copy table | Configure columns | Scatter plot | Violin plot | Export as CSV... | Showing 38/38 rows and 6/26 columns. | Copy Prompt

| Sample | STR D2 MSN | Astrocyte | STR D1 MSN | CN ST18 GABA | Oligodendrocyte | STR Hybrid MSN |
|---|---|---|---|---|---|---|
| **GSM9227428** | 696 | 1 663 | 649 | 334 | 565 | 245 |
| **GSM9227429** | 1 963 | 817 | 1 231 | 125 | 602 | 50 |
| **GSM9227430** | 808 | 1 136 | 1 642 | 170 | 921 | 59 |
| **GSM9227431** | 868 | 1 343 | 808 | 649 | 552 | 120 |
| **GSM9227432** | 1 967 | 188 | 1 201 | 152 | 742 | 218 |
| **GSM9227433** | 2 072 | 129 | 1 082 | 87 | 1 154 | 74 |
| **GSM9227434** | 1 910 | 57 | 1 248 | 123 | 560 | 515 |
| **GSM9227435** | 2 523 | 71 | 946 | 122 | 909 | 140 |
| **GSM9227436** | 1 207 | 1 754 | 500 | 170 | 314 | 108 |
| **GSM9227437** | 1 749 | 964 | 1 088 | 219 | 269 | 315 |
| **GSM9227438** | 2 432 | 513 | 1 099 | 82 | 616 | 98 |
| **GSM9227439** | 2 201 | 802 | 733 | 148 | 509 | 227 |
| **GSM9227440** | 2 687 | 448 | 936 | 124 | 337 | 103 |
| **GSM9227441** | 1 623 | 170 | 587 | 77 | 2 071 | 69 |
| **GSM9227442** | 2 528 | 674 | 714 | 94 | 268 | 92 |
| **GSM9227443** | 2 119 | 504 | 651 | 81 | 1 137 | 139 |
| **GSM9227444** | 2 383 | 348 | 643 | 52 | 1 297 | 68 |
| **GSM9227445** | 1 763 | 566 | 1 014 | 61 | 1 297 | 46 |
| **GSM9227446** | 2 153 | 347 | 815 | 69 | 1 399 | 46 |
| **GSM9227447** | 2 794 | 221 | 566 | 73 | 1 058 | 45 |
| **GSM9227448** | 2 499 | 918 | 913 | 171 | 158 | 145 |
| **GSM9227449** | 2 256 | 452 | 674 | 101 | 981 | 104 |
| **GSM9227450** | 1 826 | 216 | 577 | 50 | 1 850 | 47 |
| **GSM9227451** | 2 205 | 605 | 696 | 107 | 732 | 265 |
| **GSM9227452** | 1 489 | 1 119 | 1 057 | 153 | 642 | 267 |
| **GSM9227453** | 1 799 | 739 | 534 | 290 | 432 | 328 |
| **GSM9227454** | 2 396 | 351 | 768 | 75 | 1 115 | 151 |
| **GSM9227455** | 3 221 | 257 | 575 | 60 | 628 | 88 |
| **GSM9227456** | 2 622 | 221 | 1 450 | 65 | 440 | 41 |
| **GSM9227457** | 1 970 | 103 | 920 | 70 | 1 657 | 42 |
| **GSM9227458** | 1 537 | 139 | 922 | 67 | 2 036 | 60 |
| **GSM9227459** | 991 | 97 | 787 | 39 | 2 780 | 50 |
| **GSM9227460** | 1 521 | 1 242 | 1 436 | 67 | 208 | 83 |
| **GSM9227461** | 2 343 | 508 | 1 345 | 44 | 272 | 74 |
| **GSM9227462** | 2 918 | 385 | 689 | 51 | 540 | 69 |
| **GSM9227463** | 2 122 | 532 | 1 439 | 69 | 539 | 47 |
| **GSM9227464** | 2 891 | 288 | 883 | 56 | 469 | 38 |
| **GSM9227465** | 2 171 | 446 | 398 | 59 | 1 131 | 77 |

### Cell Type Probabilities

Average max cell type probabilities used to assign the cell types (greater is better).

Copy table | Configure columns | Scatter plot | Violin plot | Export as CSV... | Showing 38/38 rows and 6/26 columns. | Copy Prompt

| Sample | ACx MEIS2 GABA | Astrocyte | CN Cholinergic GABA | CN LAMP5 CXCL14 GABA | CN LAMP5 LHX6 GABA | CN LHX8 GABA |
|---|---|---|---|---|---|---|
| GSM9227428 | 0.2 | 0.3 | 0.3 | | 0.3 | 0.3 |
| GSM9227429 | 0.2 | 0.3 | 0.4 | | 0.3 | 0.3 |
| GSM9227430 | 0.2 | 0.3 | 0.4 | | | 0.4 |
| GSM9227431 | 0.2 | 0.2 | 0.3 | 0.3 | 0.2 | 0.3 |
| GSM9227432 | 0.2 | 0.2 | 0.3 | 0.3 | 0.2 | 0.3 |
| GSM9227433 | 0.3 | 0.2 | 0.4 | 0.3 | 0.5 | 0.3 |
| GSM9227434 | 0.3 | 0.2 | 0.4 | 0.3 | 0.2 | 0.3 |
| GSM9227435 | 0.3 | 0.3 | 0.4 | 0.2 | 0.3 | 0.2 |
| GSM9227436 | 0.2 | 0.3 | 0.3 | 0.2 | 0.2 | 0.3 |
| GSM9227437 | 0.2 | 0.2 | 0.3 | 0.3 | 0.2 | 0.3 |
| GSM9227438 | 0.2 | 0.3 | 0.4 | | | 0.2 |
| GSM9227439 | 0.2 | 0.3 | 0.3 | 0.3 | 0.3 | 0.2 |
| GSM9227440 | | 0.2 | 0.4 | | | 0.2 |
| GSM9227441 | 0.2 | 0.2 | 0.4 | | 0.3 | 0.2 |
| GSM9227442 | 0.2 | 0.3 | 0.4 | 0.1 | 0.3 | 0.2 |
| GSM9227443 | 0.2 | 0.3 | 0.4 | 0.3 | 0.2 | 0.2 |
| GSM9227444 | 0.2 | 0.2 | 0.4 | | 0.2 | 0.3 |
| GSM9227445 | 0.3 | 0.3 | 0.4 | | | 0.2 |
| GSM9227446 | 0.2 | 0.2 | 0.4 | | 0.2 | |
| GSM9227447 | 0.2 | 0.2 | 0.4 | 0.2 | 0.3 | |
| GSM9227448 | 0.2 | 0.3 | 0.4 | 0.2 | 0.2 | 0.3 |
| GSM9227449 | 0.2 | 0.2 | 0.4 | 0.2 | | 0.2 |
| GSM9227450 | 0.2 | 0.2 | 0.3 | 0.2 | 0.2 | 0.2 |
| GSM9227451 | 0.2 | 0.2 | 0.4 | 0.3 | 0.2 | 0.2 |
| GSM9227452 | 0.2 | 0.2 | 0.3 | 0.3 | 0.2 | 0.3 |
| GSM9227453 | 0.2 | 0.2 | 0.3 | 0.3 | 0.2 | 0.2 |
| GSM9227454 | 0.2 | 0.2 | 0.4 | 0.2 | 0.2 | 0.2 |
| GSM9227455 | 0.2 | 0.2 | 0.3 | 0.3 | 0.2 | 0.2 |
| GSM9227456 | 0.3 | 0.3 | 0.5 | 0.3 | | 0.2 |
| GSM9227457 | | 0.2 | 0.4 | 0.3 | 0.2 | 0.2 |
| GSM9227458 | 0.2 | 0.3 | 0.4 | 0.3 | 0.2 | 0.2 |
| GSM9227459 | | 0.3 | 0.4 | | 0.2 | 0.2 |
| GSM9227460 | 0.2 | 0.3 | 0.4 | 0.2 | 0.3 | 0.2 |
| GSM9227461 | 0.2 | 0.3 | 0.4 | 0.3 | 0.2 | 0.2 |
| GSM9227462 | 0.3 | 0.3 | 0.3 | 0.3 | 0.3 | 0.2 |
| GSM9227463 | 0.3 | 0.2 | 0.4 | 0.2 | 0.4 | 0.3 |
| GSM9227464 | 0.3 | 0.3 | 0.4 | 0.2 | 0.3 | 0.4 |
| GSM9227465 | 0.2 | 0.3 | 0.3 | 0.2 | 0.2 | 0.2 |

### Spatial Neighbors

Cell type immediate neighborhood across samples.

Percentages

STR D2 MSN | Oligodendrocyte | STR D1 MSN | Astrocyte | NA | CN ST18 GABA | Pericyte | STR Hybrid MSN | CN Cholinergic GABA | STR SST CHODL GABA | Endo | F GABA | OT Granular GABA | STR RSPO2 GABA | Lymphocyte | ACx MEIS2 GABA | VLMC | CN LHX8 GABA | OPC | SMC | CN LAMP5 CXCL14 GABA | COP | Ependymal | CN LAMP5 LHX6 GABA | Macrophage | Microglia | CN VIP GABA

Copy prompt | Export...

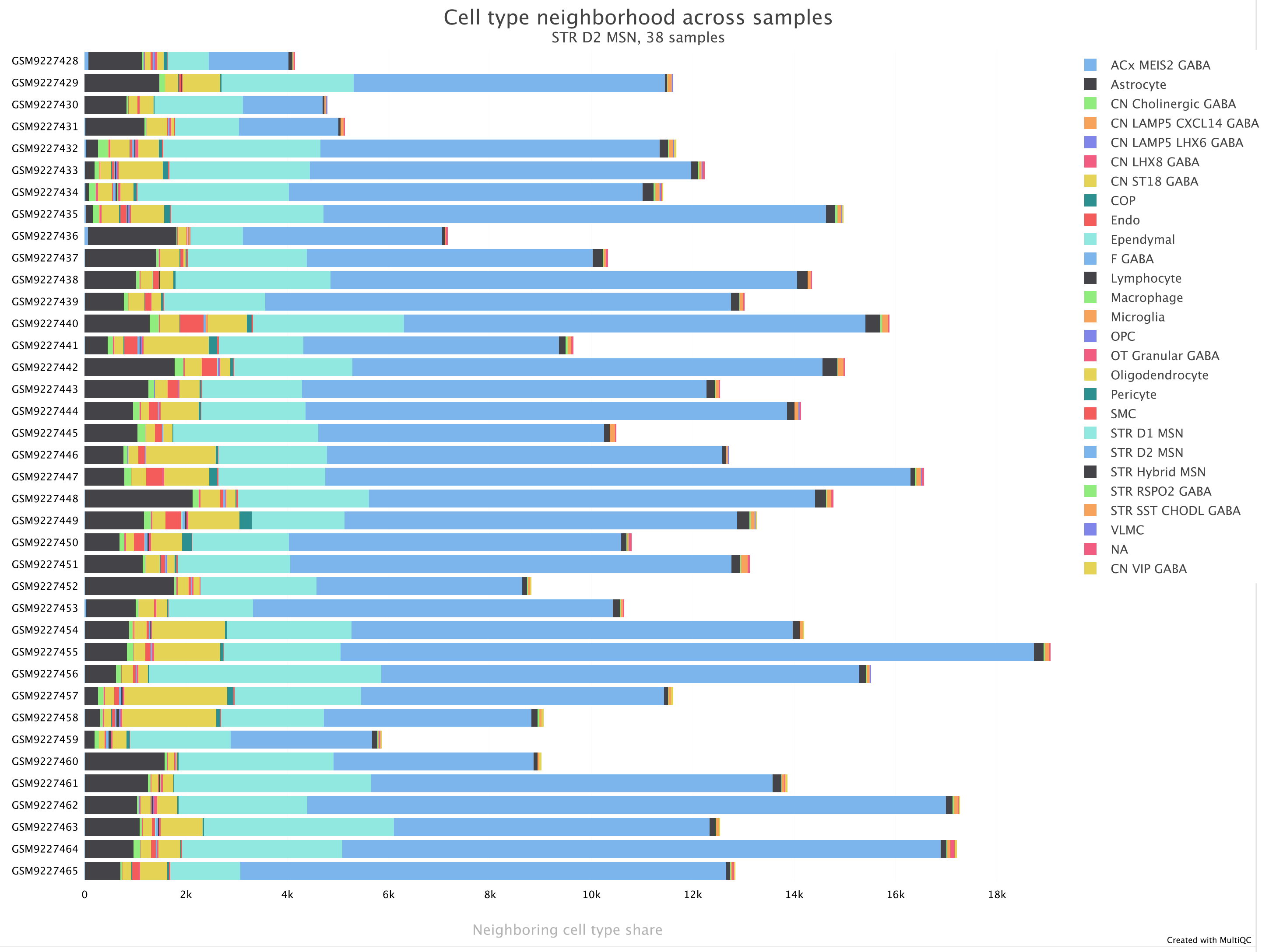

### Squidpy Ligrec Interactions

Mean interaction across samples shown as no groups were specified.

Sorted by sample | Clustered

Copy prompt | Export...

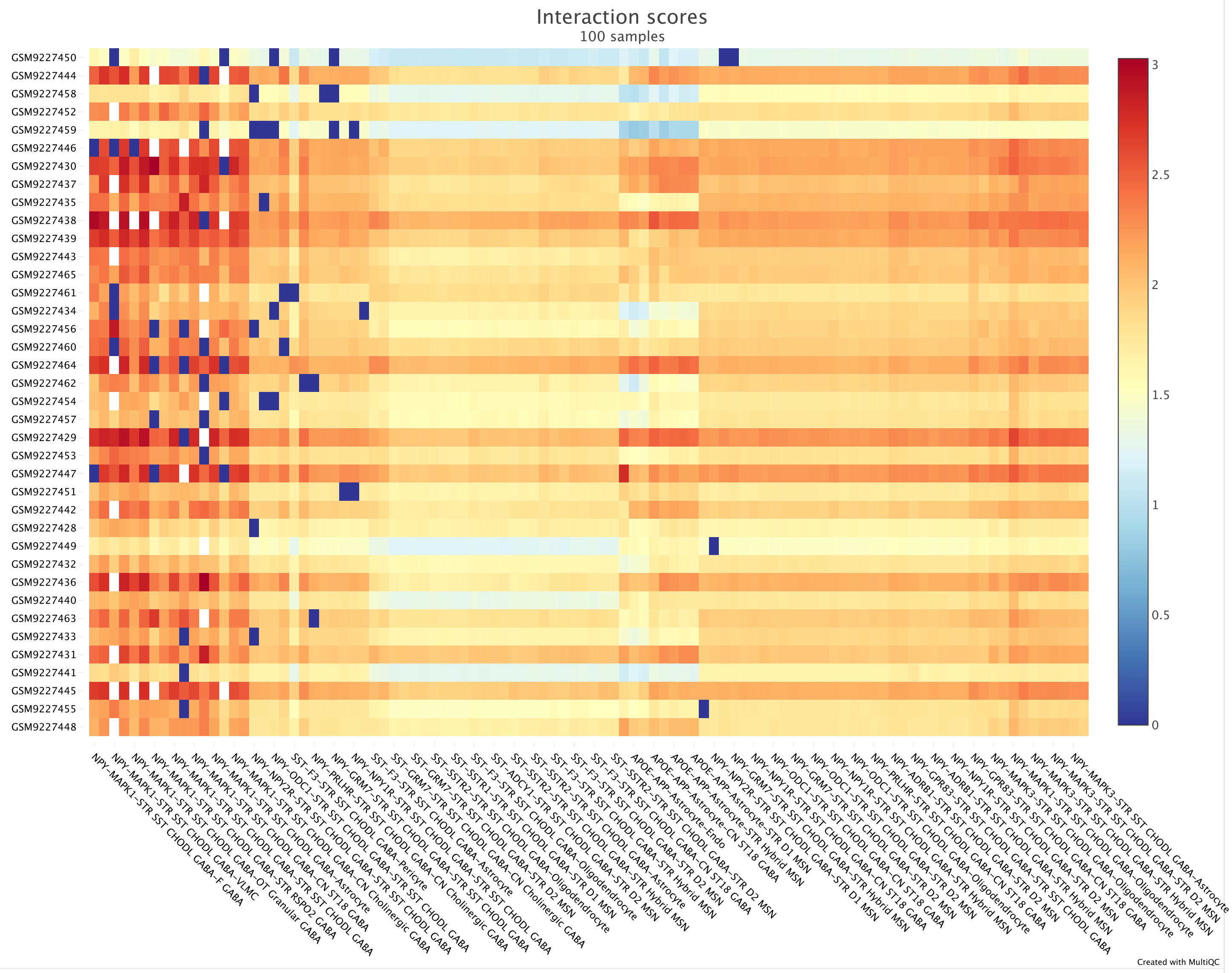

### Moran I Interactions

Mean interaction across samples shown as no groups were specified.

Sorted by sample | Clustered

Copy prompt | Export...

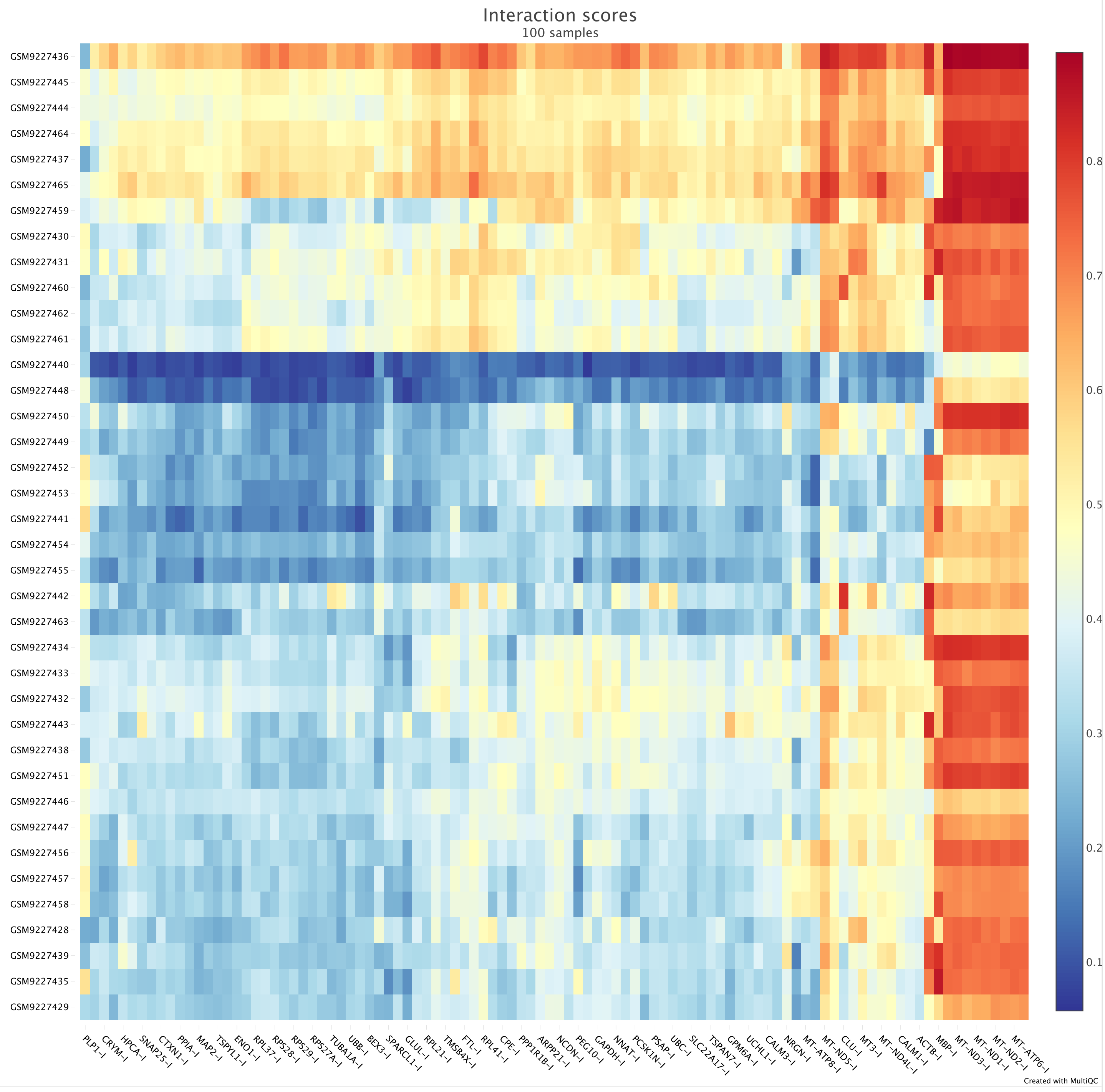

### Software Versions

Software Versions lists versions of software tools extracted from file contents.

Copy table

| Group | Software | Version |
|---|---|---|
| ADATA_ADD_METADATA | anndata | 0.12.6 |
| ADATA_FROM_VISIUM | spatialdata_io | 0.5.1 |
| | squidpy | 1.6.6.dev24+g32789aefd |
| ATLAS_GET | boto3 | 1.41.2 |
| | requests | 2.32.5 |
| ATLAS_MATCH | anndata | 0.12.6 |
| | numpy | 2.3.5 |
| QC | anndata | 0.12.6 |
| | pandas | 2.3.3 |
| RCTD | Matrix | 1.7-4 |
| | R | 4.5.2 |
| | reticulate | 1.44.1 |
| | spacexr | 2.2.1 |
| SQUIDPY_LIGREC_ANALYSIS | anndata | 0.12.6 |
| | scanpy | 1.11.5 |
| | squidpy | 1.6.6.dev24+g32789aefd |
| SQUIDPY_SPATIAL_PLOTS | anndata | 0.12.6 |
| | numpy | 2.3.5 |
| | python | 3.11.14 |
| | squidpy | 1.6.6.dev24+g32789aefd |
| STAPLE_XSAMPLE | anndata | 0.12.6 |
| | json | 2.0.9 |
| | numpy | 2.3.5 |
| | pandas | 2.3.3 |
| | scipy | 1.16.3 |
| Workflow | Nextflow | 24.10.4 |
| | STAPLE | v2.2.2-gfbf88bd |

**MultiQC v1.28** - Written by Phil Ewels, available on GitHub.
This report uses Plotly, jQuery, jQuery UI, Bootstrap and FileSaver.js.

seqera
