## Supplement 6 - NAc study workflow report for "STAPLE: automating spatial transcriptomics analysis and AI interpretation"

Nextflow Report   Summary   Resources   Tasks   [pedantic_darwin]

### Nextflow workflow report

#### [pedantic_darwin]

Workflow execution completed successfully!

**Run times**

06-Feb-2026 16:36:47 - 06-Feb-2026 18:29:51 (duration: **1h 53m 4s**)

424 succeeded

**Nextflow command**

```
nextflow run break-through-cancer/btc-spatial-pipelines -profile igs --input samplesheet.csv --outdir outs-subclass --ref_scrna_type_col
Subclass --deconvolve.n_top_genes 99999 --analyze.xsample -w /usr/local/scratch/dlvovs/work --ref_scrna /local/projects-
t3/fertig_staple/staple-paper/data/Human_HMBA_basalganglia_AIT_pre-print_NAC.h5ad
```

| | |
|---|---|
| **CPU-Hours** | 328.3 (0% failed) |
| **Launch directory** | /autofs/projects-t3/fertig_staple/staple-paper |
| **Work directory** | /usr/local/scratch/dlvovs/work |
| **Project directory** | /home/dlvovs/.nextflow/assets/break-through-cancer/btc-spatial-pipelines |
| **Script name** | main.nf |
| **Script ID** | 5a95f7d52dc1cd094f2b1163a4ea12a7 |
| **Workflow session** | 37b4bcc0-53d8-48a5-8b26-d5b1e9dd8188 |
| **Workflow repository** | https://github.com/break-through-cancer/btc-spatial-pipelines , revision main (commit hash fbf88bd923633b6b4c991a7b70b501f1908f1994 ) |
| **Workflow profile** | igs |
| **Nextflow version** | version 24.10.4, build 5934 (20-01-2025 16:47 UTC) |

### Resource Usage

These plots give an overview of the distribution of resource usage for each process.

#### CPU

Raw Usage   % Allocated

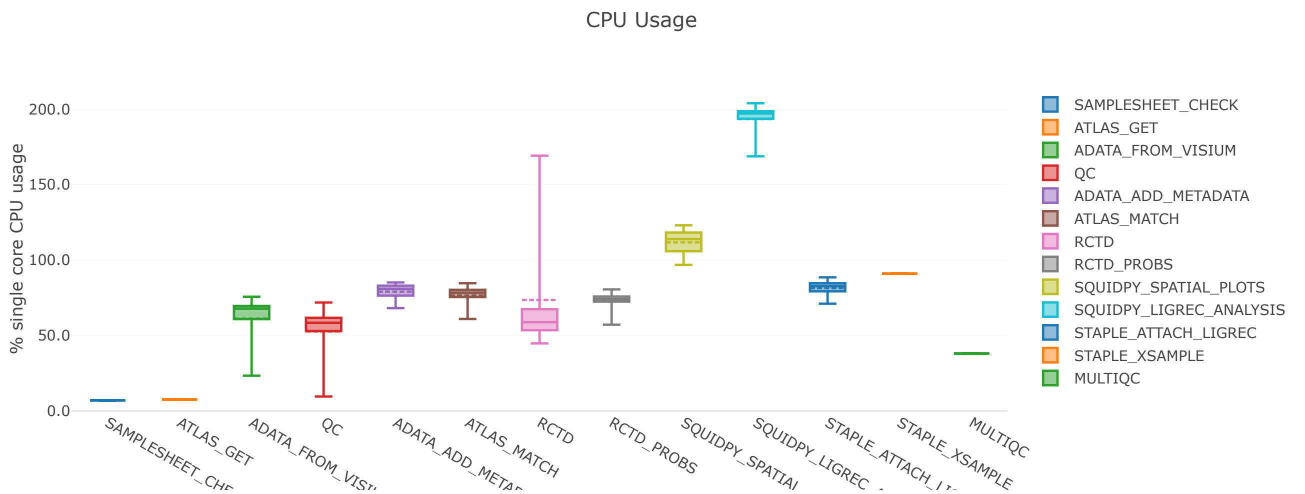

Physical (RAM) | Virtual (RAM + Disk swap) | % RAM Allocated

Physical Memory Usage

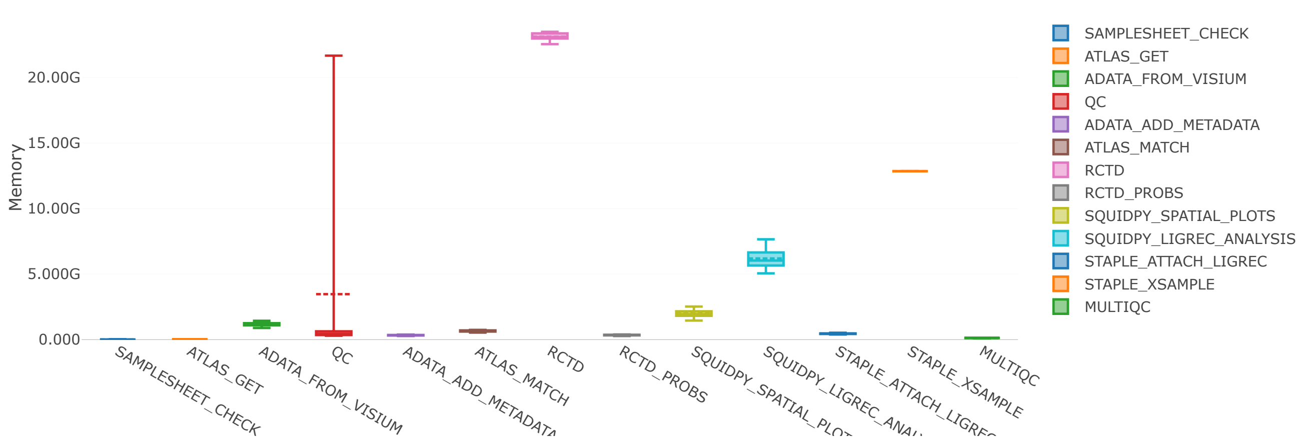


#### Job Duration

Raw Usage | % Allocated

Task execution real-time

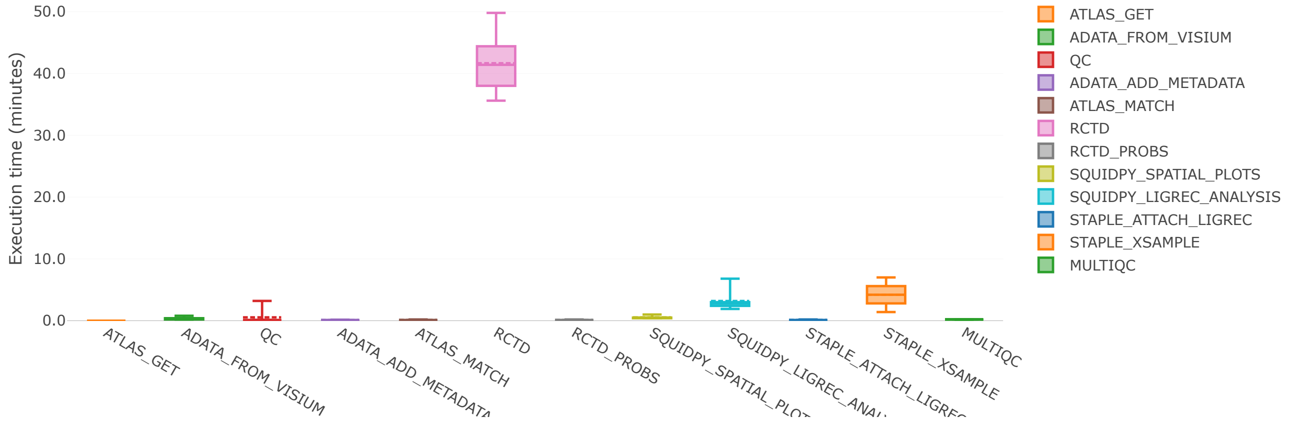


## I/O

Read | Write

Number of bytes read

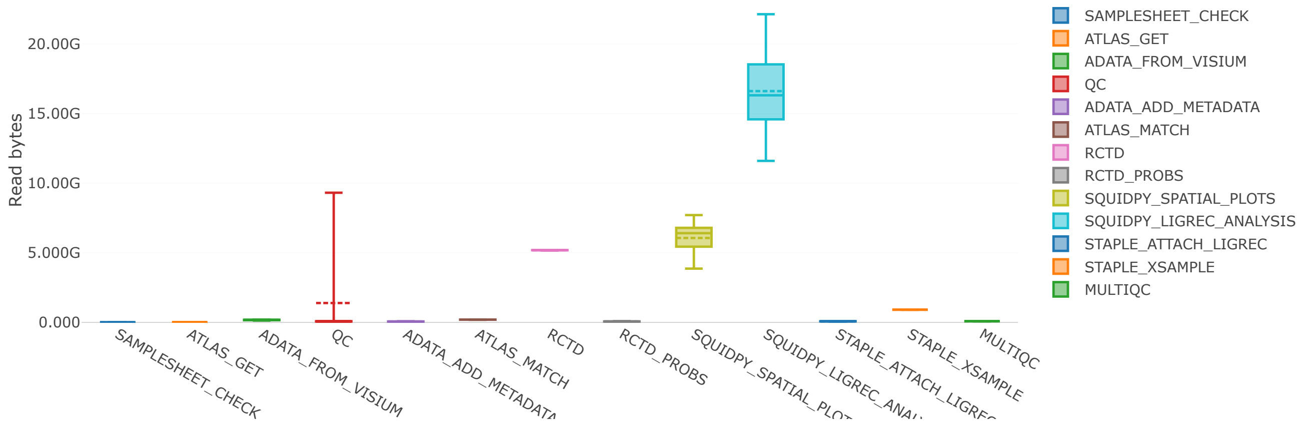

### Tasks

This table shows information about each task in the workflow. Use the search box on the right to filter rows for specific values. Clicking headers will sort the table by that value and scrolling side to side will reveal more columns.

Values shown as: Human readable

Show 25 entries Filter: Metrics Metadata All Search:

| task_id | process | tag | status | hash | allocated cpus | %cpu | allocated memory | %mem | vm |
|---|---|---|---|---|---|---|---|---|---|
| 1 | BTC:STAPLE:INPUT_CHECK:SAMPL | samplesheet.csv | COMPLETED | f3/df37fc | 1 | 7.0 | 8.000 GB | 0.0 | 11.( MB |
| 2 | BTC:STAPLE:LOAD_DATASET:ATLA | - | COMPLETED | f6/e764d0 | 2 | 7.6 | 8.000 GB | 0.0 | 28. MB |
| 3 | BTC:STAPLE:LOAD_DATASET:ADAT | - | COMPLETED | 7f/9d99b8 | 4 | 39.1 | 32.000 GB | 0.2 | 2.5 |
| 4 | BTC:STAPLE:LOAD_DATASET:ADAT | - | COMPLETED | cc/61d053 | 4 | 41.6 | 32.000 GB | 0.3 | 2.7 |
| 5 | BTC:STAPLE:LOAD_DATASET:ADAT | - | COMPLETED | ad/468878 | 4 | 24.0 | 32.000 GB | 0.1 | 2.3 |
| 6 | BTC:STAPLE:LOAD_DATASET:ADAT | - | COMPLETED | e7/471321 | 4 | 41.4 | 32.000 GB | 0.3 | 2.6 |
| 7 | BTC:STAPLE:LOAD_DATASET:ADAT | - | COMPLETED | 72/d6afd9 | 4 | 23.4 | 32.000 GB | 0.1 | 2.5 |
| 8 | BTC:STAPLE:LOAD_DATASET:ADAT | - | COMPLETED | de/37c236 | 4 | 60.4 | 32.000 GB | 0.1 | 2.4 |
| 9 | BTC:STAPLE:LOAD_DATASET:ADAT | - | COMPLETED | 47/c0b8bb | 4 | 69.6 | 32.000 GB | 0.2 | 2.5 |
| 10 | BTC:STAPLE:LOAD_DATASET:ADAT | - | COMPLETED | a5/cb314b | 4 | 68.7 | 32.000 GB | 0.2 | 2.5 |

| | | | | | | | | | |
|---|---|---|---|---|---|---|---|---|---|
| 12 | BTC:STAPLE:LOAD_DATASET:ADAT | - | COMPLETED | ae/d32c3f | 4 | 60.8 | 32.000 GB | 0.1 | 2.7 |
| 13 | BTC:STAPLE:LOAD_DATASET:ADAT | - | COMPLETED | 06/81a724 | 4 | 58.4 | 32.000 GB | 0.1 | 2.7 |
| 14 | BTC:STAPLE:LOAD_DATASET:ADAT | - | COMPLETED | 25/4404da | 4 | 69.6 | 32.000 GB | 0.1 | 2.6 |
| 15 | BTC:STAPLE:LOAD_DATASET:ADAT | - | COMPLETED | c2/b853c2 | 4 | 70.8 | 32.000 GB | 0.2 | 2.5 |
| 16 | BTC:STAPLE:LOAD_DATASET:ADAT | - | COMPLETED | 71/515dca | 4 | 69.9 | 32.000 GB | 0.2 | 2.3 |
| 17 | BTC:STAPLE:LOAD_DATASET:ADAT | - | COMPLETED | 87/6682dd | 4 | 58.3 | 32.000 GB | 0.1 | 2.1[illegible] |
| 18 | BTC:STAPLE:LOAD_DATASET:ADAT | - | COMPLETED | e0/b810f6 | 4 | 71.0 | 32.000 GB | 0.2 | 2.7 |
| 19 | BTC:STAPLE:LOAD_DATASET:ADAT | - | COMPLETED | f9/531e50 | 4 | 68.0 | 32.000 GB | 0.1 | 2.7 |
| 20 | BTC:STAPLE:LOAD_DATASET:ADAT | - | COMPLETED | 8b/bf4b71 | 4 | 57.6 | 32.000 GB | 0.1 | 2.0 |
| 21 | BTC:STAPLE:LOAD_DATASET:ADAT | - | COMPLETED | 3e/cd5e17 | 4 | 69.2 | 32.000 GB | 0.2 | 2.6 |
| 22 | BTC:STAPLE:LOAD_DATASET:ADAT | - | COMPLETED | df/6c447c | 4 | 69.7 | 32.000 GB | 0.2 | 2.7 |
| 23 | BTC:STAPLE:LOAD_DATASET:ADAT | - | COMPLETED | 77/b79c48 | 4 | 69.3 | 32.000 GB | 0.2 | 2.6 |

| | | | | | | | | | |
|---|---|---|---|---|---|---|---|---|---|
| 25 | BTC:STAPLE:LOAD_DATASET:ADAT | - | COMPLETED | 30/ceca79 | 4 | 62.2 | 32.000 GB | 0.1 | 2.2 |

Showing 1 to 25 of 425 entries

Previous 1 2 3 4 5 ... 17 Next
