## Supplement 7 - NAc study AI summary for "STAPLE: automating spatial transcriptomics analysis and AI interpretation"

Below is a structured analysis of the **MultiQC report** you attached, followed by a comparison to the **bioRxiv preprint on the human NAc cellular and spatiomolecular architecture** that you referenced.

---

### What's in your MultiQC report (high-level)

**Scope & inputs**

- **Samples**: 38 capture areas (GSM9227428–GSM9227465) from nucleus accumbens sections of neurotypical postmortem donors. Metadata include donor IDs (e.g., Br2720, Br8492), region "nucleus accumbens," and donor ages (~33–62 years). [1]
- **Atlas used for annotation**: Human_HMBA_basalganglia_AIT_pre-print_NAC (36,601 genes; **116,884 cells**; mean genes/cell ≈ 5,239). [2]

**QC / data characteristics**

- **Per-sample nuclei**: ~4,974–4,992 per capture area (constant n_cells in table). **Mean genes by counts** span roughly ~875–2,710 across samples; **mean_total_nnz_counts** ~1.6–2.6 (units as in your workflow). These ranges indicate expected variation in RNA capture across sections/slides. [3]

**Cell-type catalog (reference & outputs)**

- **Atlas cell-type composition** (reference): Dominated by **STR D1 MSN** (32,743 cells) and **STR D2 MSN** (32,219), followed by **Astrocytes** (14,136) and **Oligodendrocytes** (9,901). Other classes present at lower counts (e.g., **STR Hybrid MSN**, **OPC**, **Microglia**, inhibitory interneuron subtypes, vascular cells). [4]
- **Per-sample deconvolution / typing** (outputs):

[1] https://somumaryland-my.sharepoint.com/personal/dlvovs_som_umaryland_edu/Documents/Microsoft%20Copilot%20Chat%20Files/NAC_study_subclass_AI_prompt.txt

[2] https://somumaryland-my.sharepoint.com/personal/dlvovs_som_umaryland_edu/Documents/Microsoft%20Copilot%20Chat%20Files/NAC_study_subclass_AI_prompt.txt

[3] https://somumaryland-my.sharepoint.com/personal/dlvovs_som_umaryland_edu/Documents/Microsoft%20Copilot%20Chat%20Files/NAC_study_subclass_AI_prompt.txt

[4] https://somumaryland-my.sharepoint.com/personal/dlvovs_som_umaryland_edu/Documents/Microsoft%20Copilot%20Chat%20Files/NAC_study_subclass_AI_prompt.txt

- Across the 38 samples, **STR MSNs (D1 and D2)** dominate, with **Astrocyte** and **Oligodendrocyte** also abundant. The leading class per sample frequently alternates between **D2-enriched** and **D1-enriched** profiles.
- Illustrative examples from the "Output Cell Types" section:
  - **GSM9227430**: STR D1 MSN = 1,642; STR D2 MSN = 808; Astrocyte = 1,136 → **D1-enriched** capture area. [5]
  - **GSM9227429**: STR D2 MSN = 1,963; STR D1 MSN = 1,231; Astrocyte = 817 → **D2-enriched** capture area. [6]
  - **GSM9227442**: STR D2 MSN = 2,528; STR D1 MSN = 714; Oligodendrocyte = 268 → strong **D2** skew. [7]

- **Assignment confidence**: Average **max cell-type probabilities** per class are typically **0.2–0.4** (occasionally up to ~0.55 for some classes), which is common when mapping heterogeneous spatial spots/nuclei to fine-grained striatal subtypes. [8]

**Spatial organization (neighbors)**

- The **Spatial Neighbors** matrices show:
  - Strong **homotypic clustering** (e.g., STR D2 neighbors STR D2; STR D1 neighbors STR D1).
  - Frequent **D1–D2 co-adjacency**, consistent with mixed MSN territories (large cross-neighbor counts).

---

[5] https://somumaryland-my.sharepoint.com/personal/dlvovs_som_umaryland_edu/Documents/Microsoft%20Copilot%20Chat%20Files/NAC_study_subclass_AI_prompt.txt

[6] https://somumaryland-my.sharepoint.com/personal/dlvovs_som_umaryland_edu/Documents/Microsoft%20Copilot%20Chat%20Files/NAC_study_subclass_AI_prompt.txt

[7] https://somumaryland-my.sharepoint.com/personal/dlvovs_som_umaryland_edu/Documents/Microsoft%20Copilot%20Chat%20Files/NAC_study_subclass_AI_prompt.txt

[8] https://somumaryland-my.sharepoint.com/personal/dlvovs_som_umaryland_edu/Documents/Microsoft%20Copilot%20Chat%20Files/NAC_study_subclass_AI_prompt.txt

- Non-neuronal neighborhoods (e.g., **Astrocyte–Oligodendrocyte**) and adjacency of MSNs with glia/vascular cells are present but typically weaker than MSN-MSN homotypic signals. [9]

**Molecular interactions & spatial autocorrelation**

- **Ligand–receptor (Squidpy)**: Your report lists neuropeptide and canonical interactions (e.g., **NPY–NPY1R**, **SST–SSTR2**, and **APOE–APP**), suggesting inhibitory neuromodulatory signaling and glia–neuron crosstalk across samples. [10]
- **Moran's I** spatial autocorrelation: High spatial structure for mitochondrial and lineage/compartment genes (e.g., **MBP / PLP1** for myelination, **GFAP** for astroglia) and core neuronal transcripts—consistent with white matter and glial domains vs. gray matter MSNs. [11]

---

### How these results compare to the bioRxiv NAc study

**Reference study (scope)**

- The preprint integrated **snRNA-seq + Visium** from **10 neurotypical donors** and **38 capture areas**, producing a **spatiomolecular atlas** with **20 transcriptionally distinct cell types** and **eight spatial domains** (D1 islands; MSN_1–3; Inhibitory; Excitatory; Endothelial/Ependymal; White matter). [12]
- They emphasize **continuous spatial gradients** of **DRD1** vs **DRD2** MSNs rather than a simple core–shell dichotomy and highlight **D1 islands (DRD1-exclusive, OPRM1-enriched)** along the medial border. [13]

---

[9] https://somumaryland-my.sharepoint.com/personal/dlvovs_som_umaryland_edu/Documents/Microsoft%20Copilot%20Chat%20Files/NAC_study_subclass_AI_prompt.txt

[10] https://somumaryland-my.sharepoint.com/personal/dlvovs_som_umaryland_edu/Documents/Microsoft%20Copilot%20Chat%20Files/NAC_study_subclass_AI_prompt.txt

[11] https://somumaryland-my.sharepoint.com/personal/dlvovs_som_umaryland_edu/Documents/Microsoft%20Copilot%20Chat%20Files/NAC_study_subclass_AI_prompt.txt

[12] https://www.biorxiv.org/content/10.1101/2025.09.10.675374v1.full.pdf

[13] https://www.biorxiv.org/content/10.1101/2025.09.10.675374v1.full.pdf

- The study reports **ligand–receptor mapping** linked to neuropsychiatric risk genes and, in public summaries, highlights **opioid system** modules (**PENK/PDYN → OPRM1**) in D1 islands. [14] [15]
- Dataset registration corroborates **20 cell populations** and **8 domains**. [16]

### 1) Sampling and scale

- **Match**: Your MultiQC reflects **38 capture areas** from NAc sections—exactly the number reported for the stitched Visium reconstructions in the preprint. [17] [18]
- **Cell numbers**: Atlas dimensions in your report (**116,884 cells**) are of the same order as the preprint's retained nuclei (**~103,785**), acknowledging that reference atlases used for deconvolution can include additional cells and preprocessing differences. [19] [20]

### 2) Cell-type catalog

- **Concordant taxonomy**: Both sources feature dominant **MSN classes (DRD1/DRD2)**, with glia (**Astrocyte, Oligodendrocyte, OPC, Microglia**), **vascular** (Endothelial, Pericyte/SMC), **ependymal**, and **multiple inhibitory interneuron subtypes** (e.g., **SST**, **VIP/LAMP5**, **cholinergic**). [21] [22]

---

[14] https://www.biorxiv.org/content/10.1101/2025.09.10.675374v1.full.pdf

[15] https://www.linkedin.com/posts/prashanthi-ravichandran-96a363a0_were-excited-to-share-our-new-preprint-activity-7373437905794441219-LDUr

[16] https://www.ncbi.nlm.nih.gov/geo/query/acc.cgi?acc=GSE307587

[17] https://somumaryland-my.sharepoint.com/personal/dlvovs_som_umaryland_edu/Documents/Microsoft%20Copilot%20Chat%20Files/NAC_study_subclass_AI_prompt.txt

[18] https://www.biorxiv.org/content/10.1101/2025.09.10.675374v1.full.pdf

[19] https://somumaryland-my.sharepoint.com/personal/dlvovs_som_umaryland_edu/Documents/Microsoft%20Copilot%20Chat%20Files/NAC_study_subclass_AI_prompt.txt

[20] https://www.biorxiv.org/content/10.1101/2025.09.10.675374v1.full.pdf

[21] https://somumaryland-my.sharepoint.com/personal/dlvovs_som_umaryland_edu/Documents/Microsoft%20Copilot%20Chat%20Files/NAC_study_subclass_AI_prompt.txt

[22] https://www.biorxiv.org/content/10.1101/2025.09.10.675374v1.full.pdf

- **Subtype depth**: The preprint resolves **six MSN subtypes** (four D1, two D2) with marker-guided cross-species alignment (e.g., **RXFP1/CPNE4/TRHDE/FOXP2** in **D1 islands**; **CALB1** in matrix-like DRD1/DRD2 populations). Your MultiQC output aggregates MSN subtypes primarily at **D1 / D2 / hybrid** resolution, so it **captures major classes** but **doesn't explicitly separate** D1-island-specific DRD1 subtypes. [23] [24]

### 3) Relative abundances across capture areas

- **Preprint expectation**: **Gradients** along the mediolateral and A-P axes yield both **D1-enriched** and **D2-enriched** territories. [25]
- **Your data**: Exactly this pattern appears; e.g., **GSM9227430** is **D1-enriched** (D1 > D2), while **GSM9227429** and **GSM9227442** are **D2-dominant** (D2 > D1). That **heterogeneity across sections** is fully consistent with the **gradient model** described in the preprint. [26] [27]

### 4) Spatial architecture & neighborhoods

- **Preprint**: Defines **8 spatial domains**, with **sharp, compartmentalized DRD1 "D1 islands"** and more **gradient-like MSN_1–3** domains. **D1 islands** show **DRD1-exclusive** composition and **OPRM1-high** signatures. [28]
- **Your neighbors matrices**: Show **strong homotypic clustering** and **robust D1–D2 adjacency**, aligning with **mixed MSN fields** and **gradients**. However, because your label set does **not include a dedicated "D1 island" class**, you **cannot directly test** the preprint's **DRD1-exclusive islands** with these outputs alone. To demonstrate D1 islands in your data, you'd want a **marker-based spatial screen** (e.g., **RXFP1, CPNE4,**

---

[23] https://www.biorxiv.org/content/10.1101/2025.09.10.675374v1.full.pdf

[24] https://somumaryland-my.sharepoint.com/personal/dlvovs_som_umaryland_edu/Documents/Microsoft%20Copilot%20Chat%20Files/NAC_study_subclass_AI_prompt.txt

[25] https://www.biorxiv.org/content/10.1101/2025.09.10.675374v1.full.pdf

[26] https://somumaryland-my.sharepoint.com/personal/dlvovs_som_umaryland_edu/Documents/Microsoft%20Copilot%20Chat%20Files/NAC_study_subclass_AI_prompt.txt

[27] https://www.biorxiv.org/content/10.1101/2025.09.10.675374v1.full.pdf

[28] https://www.biorxiv.org/content/10.1101/2025.09.10.675374v1.full.pdf

**TRHDE, FOXP2, GABRQ, OPRM1**) and/or a **domain caller** (e.g., PRECAST) run on your sections; those specific diagnostics are not yet in the MultiQC tables. [29] [30]

### 5) Ligand–receptor signaling

- **Preprint**: Emphasizes domain-specific neuromodulatory circuits and **opioid-related** signaling (**PENK/PDYN → OPRM1**) within **D1 islands**; also mentions **trait-linked LR pairs** (neuropsychiatric risk genes). [31] [32]
- **Your report**: The Squidpy section surfaces **neuropeptide systems** (**NPY–NPY1R**, **SST–SSTR2**) and **glia–neuron pairs** (**APOE–APP**), consistent with **inhibitory neuromodulation** and **glial interactions**. To directly mirror the preprint's emphasis, you'd need to **ensure the LR database includes OPRM1** receptor with **PENK/PDYN** ligands and re-run the LR analysis stratified by spatial domains; those pairs aren't explicitly listed in your current tables. [33] [34]

### 6) Spatial autocorrelation (Moran's I)

- **Preprint**: Reports spatial domains for **white matter** (PLP1/MBP-rich), **endo/ependymal**, and **MSN** regions; strong spatial structure expected for lineage markers. [35]
- **Your data**: Moran's I highlights **MBP/PLP1**, **GFAP**, and core neuronal/glial transcripts as highly structured—**precisely matching** the **white-/gray-matter** and **glial** compartmentalization described in the atlas. [36]

---

[29] https://somumaryland-my.sharepoint.com/personal/dlvovs_som_umaryland_edu/Documents/Microsoft%20Copilot%20Chat%20Files/NAC_study_subclass_AI_prompt.txt

[30] https://www.biorxiv.org/content/10.1101/2025.09.10.675374v1.full.pdf

[31] https://www.biorxiv.org/content/10.1101/2025.09.10.675374v1.full.pdf

[32] https://www.linkedin.com/posts/prashanthi-ravichandran-96a363a0_were-excited-to-share-our-new-preprint-activity-7373437905794441219-LDUr

[33] https://somumaryland-my.sharepoint.com/personal/dlvovs_som_umaryland_edu/Documents/Microsoft%20Copilot%20Chat%20Files/NAC_study_subclass_AI_prompt.txt

[34] https://www.biorxiv.org/content/10.1101/2025.09.10.675374v1.full.pdf

[35] https://www.biorxiv.org/content/10.1101/2025.09.10.675374v1.full.pdf

[36] https://somumaryland-my.sharepoint.com/personal/dlvovs_som_umaryland_edu/Documents/Microsoft%20Copilot%20Chat%20Files/NAC_study_subclass_AI_prompt.txt

### Where your report **agrees** with the preprint

- **Sampling & sections**: 38 capture areas; coverage consistent with the stitched Visium reconstructions in the study. [37] [38]
- **Major classes**: Dominant **D1/D2 MSNs** plus comprehensive **glia/vascular/ependymal** and **interneuron** classes; overall taxonomy is concordant. [39] [40]
- **Gradients**: Presence of both **D1- and D2-enriched** capture areas mirrors the **gradient model** (rather than a binary core–shell split). [41] [42]
- **Spatial structure**: **Homotypic clustering** and **white matter / glial** domains with high spatial autocorrelation match domain-level organization. [43] [44]

### Gaps or partial alignment (and why)

- **D1 islands not explicitly labeled**: Your labeling schema lacks an explicit **"D1 island"** class; thus, the **DRD1-exclusive, OPRM1-high** compartments highlighted by the

[37] https://somumaryland-my.sharepoint.com/personal/dlvovs_som_umaryland_edu/Documents/Microsoft%20Copilot%20Chat%20Files/NAC_study_subclass_AI_prompt.txt

[38] https://www.biorxiv.org/content/10.1101/2025.09.10.675374v1.full.pdf

[39] https://somumaryland-my.sharepoint.com/personal/dlvovs_som_umaryland_edu/Documents/Microsoft%20Copilot%20Chat%20Files/NAC_study_subclass_AI_prompt.txt

[40] https://www.biorxiv.org/content/10.1101/2025.09.10.675374v1.full.pdf

[41] https://somumaryland-my.sharepoint.com/personal/dlvovs_som_umaryland_edu/Documents/Microsoft%20Copilot%20Chat%20Files/NAC_study_subclass_AI_prompt.txt

[42] https://www.biorxiv.org/content/10.1101/2025.09.10.675374v1.full.pdf

[43] https://somumaryland-my.sharepoint.com/personal/dlvovs_som_umaryland_edu/Documents/Microsoft%20Copilot%20Chat%20Files/NAC_study_subclass_AI_prompt.txt

[44] https://www.biorxiv.org/content/10.1101/2025.09.10.675374v1.full.pdf

preprint can't yet be **formally isolated** from your neighbors or counts tables. This is a modeling/labeling gap rather than a biological contradiction. [45] [46]

- **Ligand–receptor focus differs**: Your LR table emphasizes **NPY / SST** systems and glia–neuron interactions; the preprint's **opioid-centric** (PENK/PDYN–**OPRM1**) findings require **including those pairs** and **domain-aware** LR analysis to achieve one-to-one comparability. [47] [48]
- **Subtype resolution**: The preprint distinguishes **6 MSN subtypes** with cross-species anchors (e.g., **RXFP1/CPNE4** signatures). Your current outputs collapse to **D1/D2/Hybrid**, so **fine-grained subtype proportions** and **domain-specific enrichment** cannot be evaluated yet. [49] [50]

---

### Recommended next steps to reach "feature parity" with the preprint

1. **Add a D1-island detection pass**
   - Re-annotate spatial spots or nuclei with a **marker panel** (e.g., **DRD1, OPRM1, RXFP1/2, CPNE4, TRHDE, GABRQ, FOXP2**, and low **OPRK1**), then **segment D1-island domains**. Tools: **PRECAST** (as in the paper), or alternative domain callers; optionally enrich with **topic modeling / NMF** to recover island-specific gene programs. [51]
2. **Refine MSN subtype granularity**
   - Incorporate **six-subtype labels** (DRD1_MSN_A–D; DRD2_MSN_A–B) from the authors' signatures (Table S2) into your reference and redo deconvolution to

---

[45] https://somumaryland-my.sharepoint.com/personal/dlvovs_som_umaryland_edu/Documents/Microsoft%20Copilot%20Chat%20Files/NAC_study_subclass_AI_prompt.txt

[46] https://www.biorxiv.org/content/10.1101/2025.09.10.675374v1.full.pdf

[47] https://somumaryland-my.sharepoint.com/personal/dlvovs_som_umaryland_edu/Documents/Microsoft%20Copilot%20Chat%20Files/NAC_study_subclass_AI_prompt.txt

[48] https://www.biorxiv.org/content/10.1101/2025.09.10.675374v1.full.pdf

[49] https://www.biorxiv.org/content/10.1101/2025.09.10.675374v1.full.pdf

[50] https://somumaryland-my.sharepoint.com/personal/dlvovs_som_umaryland_edu/Documents/Microsoft%20Copilot%20Chat%20Files/NAC_study_subclass_AI_prompt.txt

[51] https://www.biorxiv.org/content/10.1101/2025.09.10.675374v1.full.pdf

quantify **domain-specific subtype enrichments** across captures (anterior/intermediate/posterior). [52]

3. **Re-run Squidpy LR with opioid and risk-gene pairs**
    - Ensure your LR database **includes PENK/PDYN→OPRM1** and other trait-linked pairs highlighted by the study; then **stratify LR inference by spatial domain** (particularly within **D1 islands**) to test the **sender/receiver maps** reported in the preprint. [53]
4. **Domain-aware neighbor analysis**
    - After calling **8 spatial domains**, recompute **neighbors** within and across domains to test **DRD1-exclusive islands** (expect reduced D2 adjacency inside islands) versus **gradient MSN domains** (expect D1–D2 adjacency). [54]
5. **Benchmark against authors' public data**
    - Cross-check your abundances and spatial signatures against **GEO GSE307587** (snRNA-seq & Visium) for donor- and position-matched sections if available. [55]

---

### Bottom line

- Your MultiQC outputs **converge strongly** with the preprint's central claims: **dominant D1/D2 MSNs**, **section-to-section gradients** in relative abundance, and **clearly structured glial/white-matter domains**. [56] [57]
- To fully recapitulate the **"cellular and spatiomolecular architecture"** emphasized by the paper—especially **D1 islands** and **opioid-related LR signaling**—you'll want to **extend the label set / domain calling** and **expand the LR panel** accordingly. [58]

---

If you'd like, I can:

---

[52] https://www.biorxiv.org/content/10.1101/2025.09.10.675374v1.full.pdf

[53] https://www.biorxiv.org/content/10.1101/2025.09.10.675374v1.full.pdf

[54] https://www.biorxiv.org/content/10.1101/2025.09.10.675374v1.full.pdf

[55] https://www.ncbi.nlm.nih.gov/geo/query/acc.cgi?acc=GSE307587

[56] https://somumaryland-my.sharepoint.com/personal/dlvovs_som_umaryland_edu/Documents/Microsoft%20Copilot%20Chat%20Files/NAC_study_subclass_AI_prompt.txt

[57] https://www.biorxiv.org/content/10.1101/2025.09.10.675374v1.full.pdf

[58] https://www.biorxiv.org/content/10.1101/2025.09.10.675374v1.full.pdf

- Parse your per-sample “Output Cell Types” and **quantify total D1 vs D2 vs glia** across all 38 sections (and compute **per-donor** summaries if you share donor–capture mappings).
- Add a **marker-based D1-island screen** on your sections and produce **domain-aware neighbor and LR** analyses.

Would you prefer I start by computing **global and donor-stratified proportions** from your report, or jump straight to a **marker-panel D1-island detection** pass?
